# Foveal vision in fast-flying birds hunting on the wing

**DOI:** 10.64898/2026.06.05.730304

**Authors:** Tania Rodrigues, Michel Marc Matter, Alain Chiodini, Bernard Genton, Emilie Brethaut, Florence Chiodini, Lidia Matter-Sadzinski, Jean-Marc Matter

## Abstract

Processing rapid motion while maintaining high spatial acuity is a fundamental evolutionary challenge for the vertebrate visual system. Here, we investigated the structural adaptations enabling aerial insectivores - swifts (*Apus apus*) and swallows (*Hirundo rustica*, *Delichon urbicum*) - to track and capture prey at high speeds. We show that these phylogenetically distinct species share highly specialized temporal foveae that provide sharp frontal vision. Strikingly, this avian specialization converges on primate foveal architecture, featuring cones with long axons and a unique cluster of large, orthotopic ganglion cells (soma area ≥ 200 µm²) surrounding a deep foveal pit. By tracking their large axons, we mapped their neural representation within the optic nerve and tectum. Despite the low abundance of these foveal cells, their substantial tectal magnification reflects high processing demands. This cluster of putative motion-sensitive ganglion cells suggests that foveal neural circuitry links high-acuity vision to rapid temporal processing in these birds.

## INTRODUCTION

Foveal vision can be regarded as the pinnacle of eyesight development in vertebrates, due to its provision of the sharpest and most detailed color vision. That level of sophistication is reached by primates and by birds of the Neoaves clade. Well over half of the roughly 10’500 bird species use foveal vision. The fovea cannot, therefore, be considered a niche-specific adaptation (Masland and Martin, 2007). As noted by Chen et al. (2019), studies of foveal visual mechanisms have become increasingly rare in the past 3 or 4 decades. This is largely due to inherent challenges in vision and primate research, which are compounded by the prevalence of mouse models. Domestic Rock Pigeon (*Columba livia domestica*) was a long-standing model for studying foveal vision in birds, but the number of studies on pigeon visual system has decreased steadily over the past 25 years, replaced by Domestic Chicken (*Gallus gallus domesticus*) as the preferred model for avian research. Unlike pigeons, chickens do not have a fovea and are not known for agile and sustained flight. While the fovea’s role in high visual acuity has long been established, whether it detects motion remains a subject of ongoing debate in visual neuroscience. We have a fairly good understanding of how birds detect global image motion, as produced by the bird’s flight, thanks to studies carried out on pigeons (Brecha and Karten, 1979; Brecha et al., 1980; Karten et al., 1977), and more recently on hummingbirds (*Calypte anna*) and zebra finches (*Taeniopygia sp.*) (Gutierrez-Ibanez et al., 2025; Smyth et al., 2022). Yet, it is not known whether birds have neurons in their retinas that handle the complex distinction between object motion and self-induced image motion. Retinal ganglion cells (GCs) that achieve this task are termed *object motion sensitive* (OMS) and have been discovered in salamanders, rabbits and mice (Baccus et al., 2008; Kim and Kerschensteiner, 2017; Kuhn and Gollisch, 2016; Olveczky et al., 2003), but not in birds. The implementation of multielectrode systems for recording light-driven activity in avian retinal cells (Seifert et al., 2023) should accelerate our understanding of avian retinal electrophysiology, which is lagging behind other species. Nonetheless, cells with interesting electrophysiological profiles would still have to be characterized at the morphological and molecular levels.

Here, we made the simple assumption that putative GCs that detect individual moving objects might be more numerous and/or noticeable in fast-flying birds feeding on insects caught in flight than in other birds. While swifts (Apodiformes) and swallows (Passeriformes) are phylogenetically distinct, they represent a striking case of convergent evolution, occupying a highly specialized, similar ecological niche. Despite this, we found that their retinal organizations differ significantly, with a small temporal area encompassing a foveal pit being their only common trait. These analogous temporal foveae displayed specialized features that we did not find in other birds that hunt prey. One of the most striking characteristics was a population of big and orthotopic GCs with soma area ≥ 200 μm^2^ that clustered around a deep temporal pit. We took advantage of the fact that the large axons originating from the few hundred big foveal GCs can be traced easily to analyze the distribution of foveal input in the optic nerve and in the midbrain. We show that they are distributed in broad but precisely delimited areas in the optic nerve. All large axons originating from the big GCs of the temporal fovea innervate the optic tectum (homologous to the superior colliculus in mammals). This enabled us to directly visualize, for the first time in vertebrates, the magnified neural representation of a fovea within the brain. We show that this overrepresentation does not result simply from the high number of foveal GCs. Instead, the disproportionate allocation of tectal tissue to foveal GCs may stem from their specialized role in recruiting more tectal neurons to process visual motion input compared to extra-foveal GCs.

## RESULTS

### 1. Swifts and swallows are highly efficient at identifying and catching insects in flight

The eye position in diurnal birds like swifts and swallows indicates a combination of frontal and lateral vision (Slonaker, 1897; Figure 1A). One behavioral trait that distinguishes the swifts and swallows from other diurnal insect-hunting birds or birds of prey is their ability to catch insects on the wing using their beaks (Figure 1B). One of us (BG) hosts a colony of Common Swifts (*Apus apus*) and a colony of House Martin (*Delichon urbicum*) at his home, providing opportunities to monitor their behaviors during the breeding season (Genton and Jacquat, 2014). Swifts form small, rounded lump of insects in their throat called bolus. These boluses are regurgitated and fed to their chicks. On rare occasions, the parents lose their bolus before delivery to their offspring, thereby providing the opportunity to analyze their content (Figure S1). Swifts distinguish between different insects, specifically avoiding stinging ones like bees and wasps. However, they readily consume harmless syrphid flies, even though these flies mimic bees and wasps to avoid predators (Figure S1; Gory, 2008). Identifying traits might include visual cues related to flight behavior, body shape, and wing venation. Notably, Syrphidae possess a single pair of wings, distinguishing them from bees and wasps, which have two pairs. Analyses of bolus and dropping samples demonstrate that swifts actively distinguish insects by size and morphology, increasing the average prey size to accommodate growing chicks (Cucco et al., 1993). However, in light of these results, it remained unconfirmed whether swifts utilize color for prey discrimination. Bolus analysis reveals that swifts heavily prey on winged black garden ants during their nuptial flight (Figure S1), while excluding winged yellow meadow ants (*Lasius flavus*), despite their high abundance near the colony. Swifts likely use color to differentiate between similar-looking yellow and black ants to avoid the former’s biting defenses.

**FIGURE 1.**
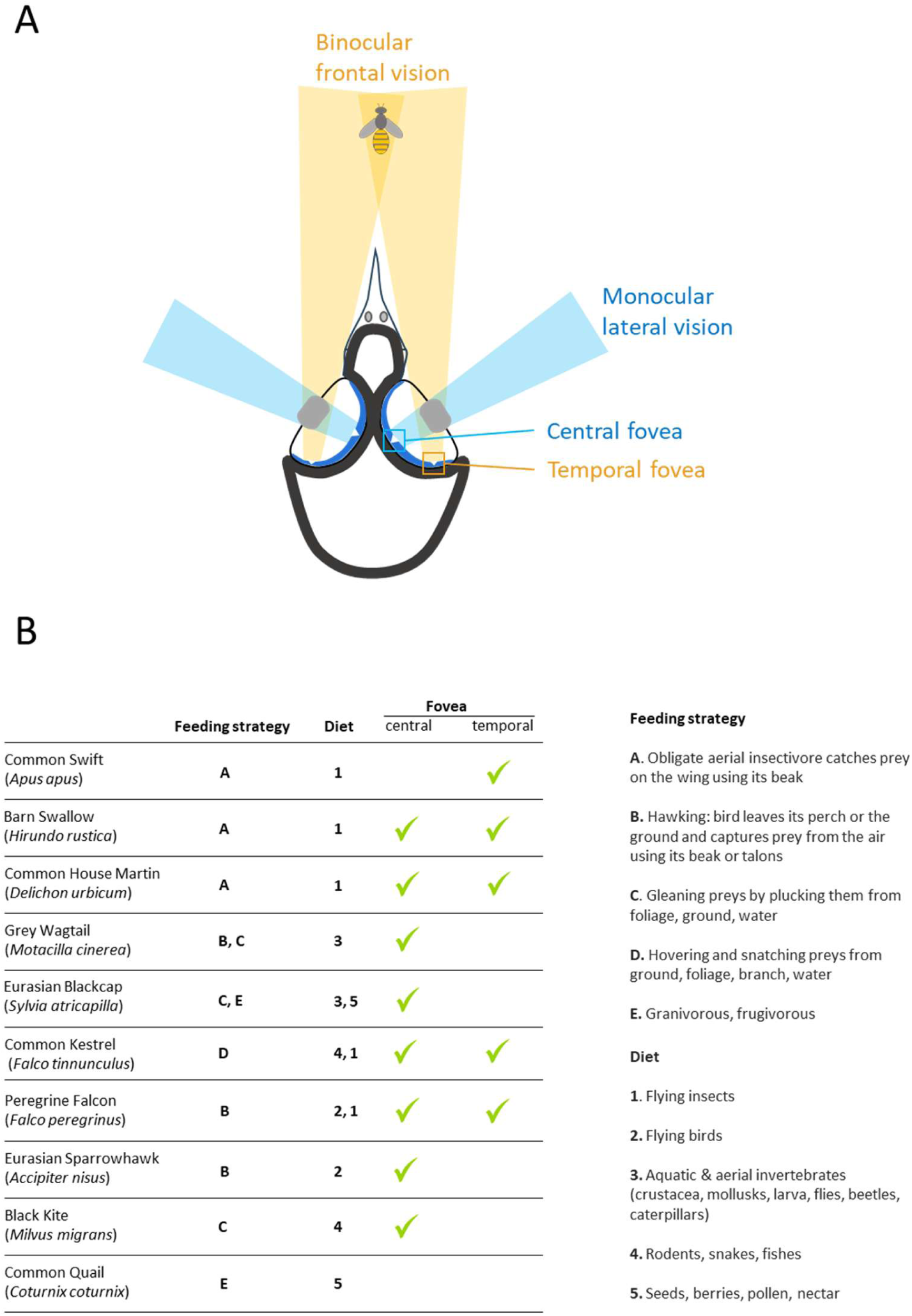
Bird foraging behavior and foveal vision. (**A**) Horizontal section of a swallow’s head and eyes showing the central and temporal foveae, with a schematic representation of frontal and lateral visual fields (adapted from Slonaker, 1897). (**B**) Hunting strategies and diets of foveate and non-foveate species investigated in this study (Table S1). While birds are opportunistic, the diets indicated represent the most common prey for each species. Notably, Common Kestrels and Peregrine Falcons are excluded from Category A. Although they occasionally hunt insects in flight, they rely on their talons rather than the bill-based capture strategy characteristic of swifts and swallows.

**FIGURE S1 (related to FIGURE 1).**
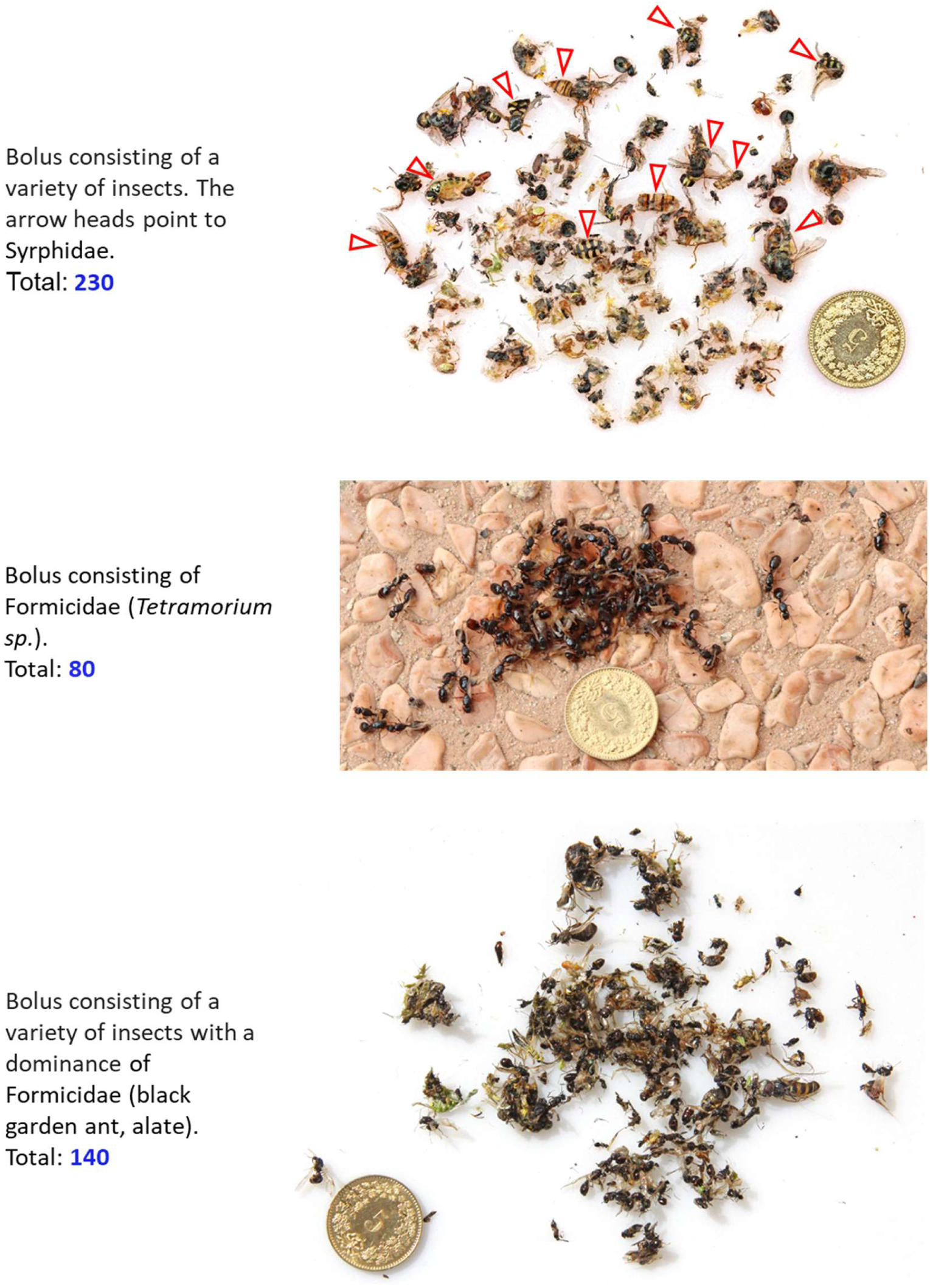
Composition and size of Common Swift boluses. Three boluses collected from a nesting site (BG’s home), illustrating typical prey clusters. While boluses generally contain a diverse range of flying insects, monospecific compositions are frequently observed. The number of individual insects identified per bolus is noted. Scale bar: 17.15 mm coin for reference.

**TABLE S1 (related to FIGURE 1).** Characteristics of the specimens analyzed in this study.

| Species | Specimen | Date of collection | Age | Sex | Injury | Duration of stay | Parameters |  |  |
| --- | --- | --- | --- | --- | --- | --- | --- | --- | --- |
|  |  |  |  |  |  |  | Body weight (g) | Brain weight (g) | Eye diameter (mm) |
| Barn Swallow<br>(*Hirundo rustica*) | Hr08 | 2020.06.15 | J | F | BF | 4d | 15.28 | 0.64 | 9 |
|  | Hr11 | 2020.08.03 | J | M | BF | 1d | 15.06 | 0.64 | 9 |
|  | Hr21 | 2021.09.14 | J | n/a | BF | 0d | 14.44 | 0.63 | 9.5 |
|  | Hr28 | 2013.08.15 | A | n/a | BF | 0d | n/a | n/a | 9.5 |
| House Martin<br>(*Delichon urbicum*) | Du15 | 2020.09.16 | J | n/a | BF | 1d | 9.61 | 0.52 | 8 |
|  | Du25 | 2021.12.06 | J | M | FP | 5m | 17.33 | 0.49 | 9 |
|  | Du26 | 2021.12.06 | J | n/a | FP | 5m | 15.96 | 0.45 | 9 |
| Common Swift<br>(*Apus apus*) | Aa06 | 2020.05.28 | A | M | FP | 0d | 32 | 0.86 | 12 |
|  | Aa13 | 2020.09.16 | J | n/a | BF | 1m | 39.66 | 0.68 | 11 |
|  | Aa14 | 2020.09.16 | J | M | FP | 1m | 33.26 | 0.69 | 11.5 |
|  | Aa18 | 2021.07.29 | J | M | FP | 3d | 35.4 | n/a | n/a |
|  | Aa19 | 2021.07.29 | J | n/a | FP | 3d | 31.86 | 0.69 | 11.5 |
| Common Kestrel<br>(*Falco tinnunculus*) | Ft03 | 2020.02.10 | A | M | BF | 9m | n/a | n/a | 20 |
|  | Ft10 | 2020.07.27 | J | n/a | BF | 16d | 158.53 | 3.8 | 18 |
| Peregrine Falcon<br>(*Falco peregrinus*) | Fp23 | 2021.10.21 | A | F | BF | 4m | 596 | 6.13 | 22 |
| Black Kite<br>(*Milvus migrans*) | Mm04 | 2020.03.12 | A | M | FP | 7m | 807 | 7.51 | 24 |
| Eurasian Sparrowhawk<br>(*Accipiter nisus*) | An01 | 2020.02.05 | A | n/a | BF | 1d | n/a | n/a | 19 |
| Grey Wagtail<br>(*Motacilla cinerea*) | Mc12 | 2020.08.17 | J | M | BF | 2d | 15.73 | 0.65 | 8 |
| Eurasian Blackcap<br>(*Sylvia atricapilla*) | Sa09 | 2020.07.22 | J | M | n/a | 1d | 11.7 | 0.8 | 8 |
| Japanese Quail<br>(*Coturnix japonica*) | Cj22 | 2021.10.18 | A | M | n/a | n/a | 324 | 0.86 | 11 |

Abbreviations: M, male; F, female; BF, bone fracture; FP, feather problem; d, day; m, month

Observing the swift colony (50 couples) during the period parents feed their young, permit to count a daily average of 25 sorties with occasional peaks reaching 40 sorties. This suggests that one bird is capable of accumulating a bolus in a period of ∼30 minutes. In the boluses, we counted between 100 and 200 insects (Figure S1), although boluses containing up to 800 insects have been recorded (Engeler, 2025). So, a swift typically captures one insect every 10 to 20 seconds while flying at speeds between 40 and 60 km/h. Despite the fact that errors in prey selection may occasionally occur (Genton and Jacquat, 2014), the success rate is breathtaking. House Martins and Barn Swallows (*Hirundo rustica*) have shorter foraging sorties compared to swifts, resulting in smaller bolus or no bolus at all. Unlike Common Swifts, House Martins frequently catch prey nearby and return immediately to feed their chicks. Swifts and swallows often share the same airspace, though swifts are known for their high-altitude, high-speed flights in vast aerial spaces, while swallows tend to fly closer to the ground or water in a more cluttered airspace where prey are less dispersed. House Martins are known for their gliding and slower movements, while Barn Swallows fly with a more rapid, arrow-like style. Among birds of prey, the Peregrine Falcon (*Falco peregrinus*) and the Eurasian Sparrowhawk (*Accipiter nisus*) (Figure 1B) are highly skilled at catching birds in flight. However, while they are known for their impressive hunting speed, their success rate is not as high as one might expect. The Peregrine Falcon hunting success rates range from 9% to 35% depending on experience, environment and weather (Dekker, 2009). Eurasian Sparrowhawks have also a low overall hunting success rate, estimated at around 10%, yet they manage to catch a few prey items daily (Cresswell et al., 2010). The Grey Wagtail (*Motacilla cinerea*) forages terrestrially along watercourses for aquatic invertebrates and make agile aerial sallies (hawking) from a perch or the ground to catch flying insects. Conversely, the Eurasian Blackcap (*Sylvia atricapilla*) is an arboreal gleaner of insects and spiders, transitioning to a fruit-based diet during autumn and winter (Figure 1B). In sum, swifts and swallows are the undisputed virtuosos of the sky, snatching prey mid-air with surgical precision. We asked what visual system adaptations enabled them to develop a distinctive lifestyle within the avian world.

### 2. The swift and swallow central retinas are different

The fact that swallows possess a central fovea while swifts do not is intriguing. Rochon-Duvigneaud (1943) and Oehme (1962) duly noted the absence of a central pit in the Common Swift (Figure 2A), but since then no one has looked at this fascinating case. Although the Common Swift can be considered as an ‘*exception*’ (Bringmann, 2019), it remains that the fovea centralis is an established tenet of high visual acuity and its absence raises a number of questions. In the swift central retina, the number and density of cells in the ganglion cell layer (GCL) were the lowest measured in birds (Figure 2B, C; Rodrigues et al., 2023). The number of cell bodies located in the inner region of the inner nuclear layer (INL) was comparatively high and we wondered whether the INL would not contain a larger than expected population of displaced ganglion cells (DGCs; Karten et al., 1977; Wylie et al., 2014). These neurons are involved in the detection of self-induced image motion, or optic flow, an essential feature of aerial birds (Gutierrez-Ibanez et al., 2025). In the GCL of the Common Swift (specimens Aa06, Aa13, Aa14, Aa19), cell bodies were distributed in a single, uniform layer over the entire surface of the retina (except for a small temporal area). It was therefore possible to obtain fairly precise total cell counts, estimated at 2.80 × 10^5^ somas (Aa14; Figure S2A). Counting the number of axons in the optic nerve (ON) provides a more accurate estimate of GC number than other methods. As presented later in this Results section, 2.78 × 10^5^ axons were counted in the ON of the specimen Aa14, suggesting that almost all GCs were in the GCL, and nearly all cells in the GCL were GCs. The highly regular distribution of neurons in the central retina enabled us to propose that 32 cones distributed over an area of ∼100 μm^2^ converge on a single GC via 24 BCs and 16 ACs (Figure 2B). Such a circuit would be well suited for detecting distant moving objects.

**FIGURE 2.**
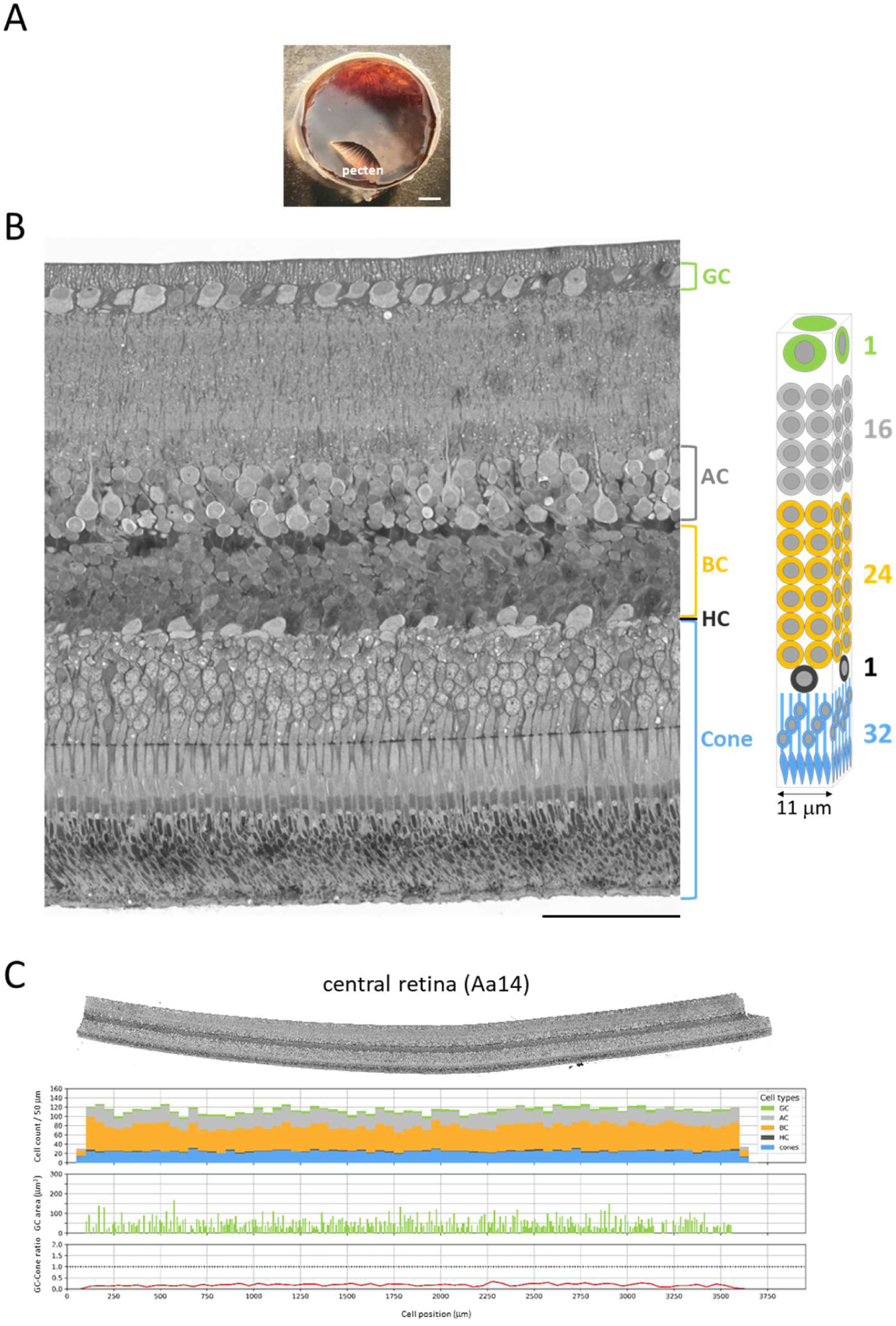
The Common Swift central retina. (**A**) Posterior eye cup. There is no foveal pit in the area centralis. (**B**) Histological analysis of the central retina, visualized on 1 µm semi-thin plastic sections stained with toluidine blue, Cones are the sole photoreceptor type within the area centralis. The highly regular distribution of the five neuron types suggests a fundamental radial module comprising 1 GC, 16 ACs, 24 BCs, 1 HC, and 32 cones. (**C**) A stacked bar chart illustrates the distribution of five retinal neuron types across a 3.5 mm tissue section, binned at 50 µm intervals. Each bar’s segments quantify the neuron types within that interval. Separate graphics present GC soma sizes and GC to cone ratios. GC, retinal ganglion cell; AC, amacrine cell; BC, bipolar cell; HC, horizontal cell; cone, cone photoreceptor. Bar scale: 2 mm (**A**). 50 μm (**B**).

In swallows, the central retinas were radically different. Among the 3 specimens of Barn Swallows (Hr08, Hr11, Hr21) and the 3 specimens of House Martins (Du15, Du25, Du26) that we have analyzed, all had a deep central pit, except for one juvenile Barn Swallow (Hr08) that had a *dome-shaped fovea* (Figure 3A). The fact that it was injured in the wing early during the breeding season (Table S1) has suggested that the lateral displacement of GCs, ACs and BCs leading to pit formation had not happened yet in that bird. The small number of specimens examined does not allow us to exclude, however, the possibility that a *dome-shaped* fovea could persist in adult. Anyway, this specimen provided us with the rare opportunity to map the distribution of foveal neurons before they move laterally (Figure 3B). Indeed, when foveal neurons become intermingled with parafoveal neurons, it makes it almost impossible to distinguish and count them. In the domed fovea, radial columns of neurons were regularly aligned. On a cross section at the maximum width of the dome (368 μm), there were 874 BCs distributed in 66 radial columns of 13±1 somata, 137 GCs distributed in 44 radial columns of 3 somata (rarely 2 or 4), 247 ACs, 63 HCs and 393 cone nuclei. Our assumption that the foveal GCL contained only GCs was based on the fact that this layer consisted of tightly packed radial columns, a cell arrangement that we did not observe elsewhere in the retina (except in the temporal fovea). Cone outer segments (OS) were long and thin (1 μm diameter) and their nuclei were stack in 6 sublayers. Similar neuron densities and neuron type distribution were found on 50 semi-thin sections encompassing the entire thickness of the dome (∼400 μm), reflecting the highly regular alignment of radial columns over a round-shaped central area of 10^5^ μm^2^. The basic module in this area would consist of 324 cones distributed on an area of ∼300 μm^2^ that converge on 12 GCs (organized in 4 radial columns of 3 GCs) through 117 BCs and 20 ACs (Figure 3B). This central fovea is characterized by a high density of cones and tightly packed GCs. However, we observe an unprecedented situation where foveal cones have a relatively high convergence ratio, making midget-like pathways rare or absent. This suggest that this fovea is well adapted for detecting details at high temporal resolution.

**FIGURE 3.**
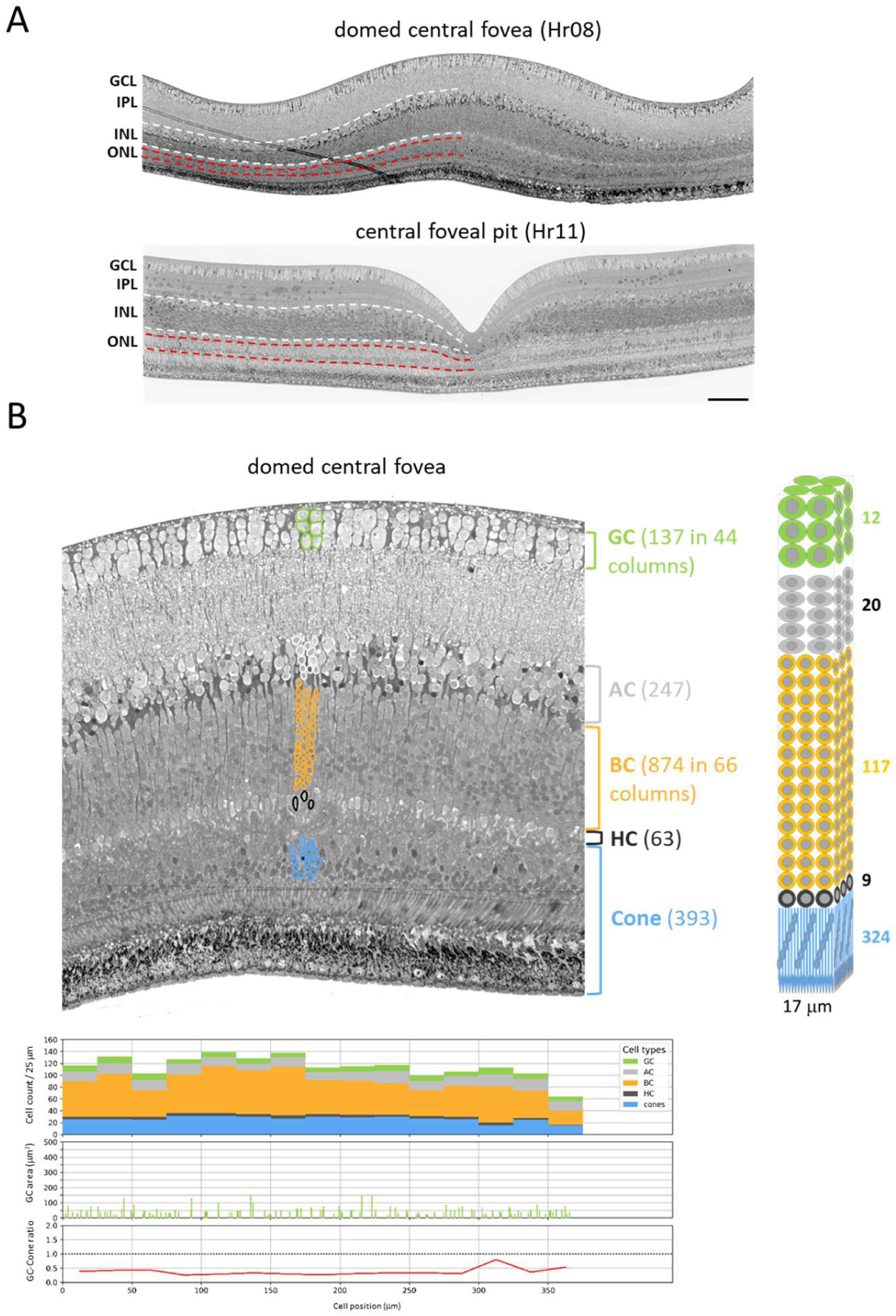
The Barn Swallow central retina. (**A**) The central retina in the specimen Hr08 (upper panel) and Hr11 (lower panel). White and red dashed lines delimit the inner nuclear layer (INL) and outer nuclear layer (ONL), respectively. Variations in their thickness highlight cytoarchitectonic differences between dome-shaped and pit-shaped foveae. (**B**) Retinal histology of the domed fovea (Hr08). Upper left panel: representative image of the fovea across its entire width. GCs and BCs are organized into distinct radial columns containing 3 and 13 somas, respectively. Each BC column is associated with a single HC. Lower left panel: Each bar is subdivided to show the distribution of the five neuron types across 25 µm segments. Data for GC soma area (µm²) and the ratio of GCs to cones are included. Right panel: The highly regular distribution of the five neuron types throughout the foveal volume suggests a fundamental radial module. Each module would comprise 4 columns of GCs, 4 columns of ACs, 9 columns of BCs, 9 HCs, and 324 cones. Based on this architecture, the fovea consists of approximately 480 modules arranged in a 22 x 22 matrix. Bar scale: 100 μm (A).

### 3. Topographic anisotropy in the Barn Swallow retina: the role of displaced amacrine cells

We wondered whether differences between the swift and swallow retinas extended beyond the central area. Cell counts along the dorso-ventral axis revealed indeed marked interspecies variations in cell distribution (Figure S2A-C). In the Common Swift, the thickness of the retina and the densities of neurons were quite uniform over the whole retinal surface, whereas in the Barn Swallow, the retina was thicker and neuron densities were higher in a broad central area compared to the periphery. In the swallows (Hr11, Hr21), the total number of cells in the GCL was estimated at 4.5 × 10^5^, i.e., 1.7 × 10^5^ more cells than in the GCL of the Common Swift. However, as presented later in this Results section, the total number of axons in the ON was smaller in the Barn Swallow than in the Common Swift (2.35 × 10^5^ in Hr21 vs. 2.78 × 10^5^ in Aa14). Intrigued by an apparent contradiction, we identified a population of small cell bodies (≤ 20 μm^2^) that formed 31% of the GCL in the Barn Swallow, whereas they were absent in the Common Swift (Figure S2D). The numbers of small axons (0.1-0.5 μm^2^) were almost the same in the optic nerves of the two species, suggesting that these 31% extra cells in the Barn Swallow’s GCL were not GC, but rather displaced amacrine cells (DAC). Many GCL cells with soma areas of 20–30 µm² could also be DACs. The proportion of DACs in birds can vary widely (Binggeli and Paule, 1969; Chen and Naito, 1999; Ehrlich, 1981; Haverkamp et al., 2021). The fact that DACs were few in number or perhaps even absent in the Common Swift does not affect this aerial bird from exhibiting flight and hunting behaviors very similar to those of the Barn Swallow. The many differences in the retina between the two species appear to contradict the widely held view of the evolution of eyes as being ‘task-led’.

**FIGURE S2 (related to FIGURES 2 and 3).**
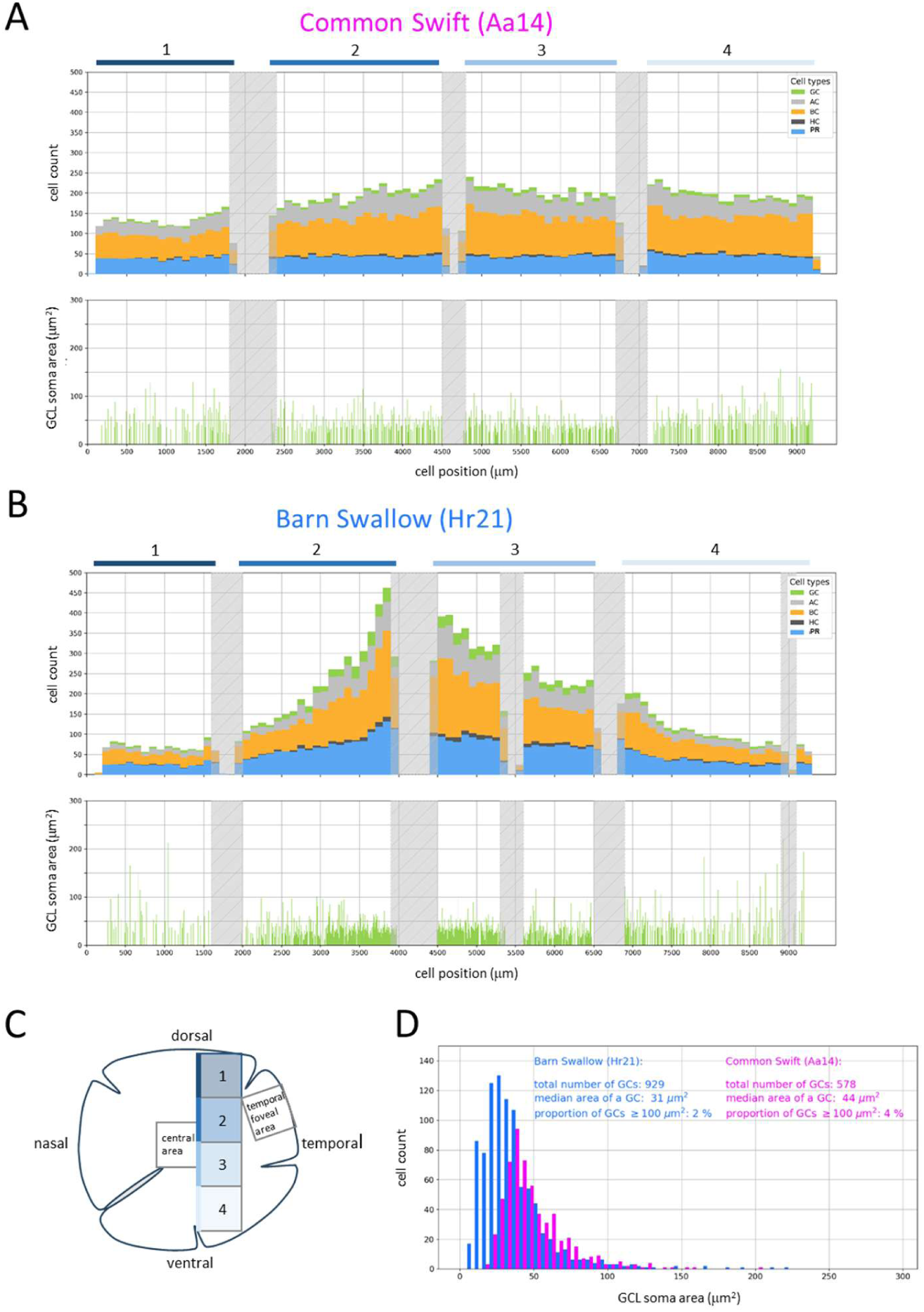
Comparative retinal histology and neuronal distribution along the dorso-ventral axis. (**A**, **B**) Retinal sections of the Common Swift (Aa14) and Barn Swallow (Hr21) divided into four adjacent sectors (1–4) spanning the ventral-to-dorsal poles (**C**). Stacked bar charts display the density of five retinal neuron types along the ∼9 mm tissue length (100 µm bins). Shaded regions indicate sector discontinuities; the shaded zone in the Barn Swallow sector 3 denotes a processing-induced fracture. (**A**, lower) In Common Swifts, somas of the Ganglion Cell Layer (GCL) form a uniform monolayer across the entire retina (except within the temporal fovea) with a total population estimated at ∼2.80 × 10⁵ cells. (**B**, lower) In the Barn Swallow GCL, somas are at the highest density in sectors 2 and 3. Across the entire retina, soma abundance is ∼1.6 times higher than in Common Swifts. The total population is estimated at ∼4.5 × 10⁵ cells. (**D**) An increased number of small somas (< 30 µm²) underlies the higher overall cell abundance in the Barn Swallow GCL.

### 4. Analogous temporal foveae

Our finding of analogous foveae in the temporal retinas of the Common Swift, the Barn Swallow and the House Martin partly resolved the above-mentioned paradox (Figures 4, S3). In the three species, a deep pit delimited a foveal area facing the center which was markedly different from the one facing the periphery. Decreased densities of cones and BCs in the foveal area oriented towards the periphery resulted from increased thickness of cone inner (IS) and outer (OS) segments (Figure 5A), and also because BCs and cone nuclei moved away from the foveal pit resulting in elongated axon connections between the IS and cone nuclei, as well as to synapses (Figure 5B). Another notable change that occurred within the foveal pit was the abrupt transition from long to short cone OS. OS were anchored to the foveolar retinal pigment epithelium (RPE), and their change in length correlated with a change in the thickness of the RPE (Figure 5A). The pit formed at the exact location where cone types changed. What particularly caught our attention were clusters of big somas (≥ 200 μm^2^) particularly abundant in the GCL of the foveal area oriented towards the periphery (Figures 4). Swifts and swallows developed analogous specialized temporal areas, despite the many differences in the organization of their retinas. Homoplasy occurring within a small retinal area is unprecedented and challenges our current understanding of how the retina develops. The idea that the temporal fovea might develop with a degree of autonomy with respect to other retinal areas is supported by the fact that the retina of the juvenile Barn Swallow Hr08 displayed a fully developed temporal fovea (Figure 5A), while its central fovea was yet dome-shaped (Figure 3).

**FIGURE 4.**
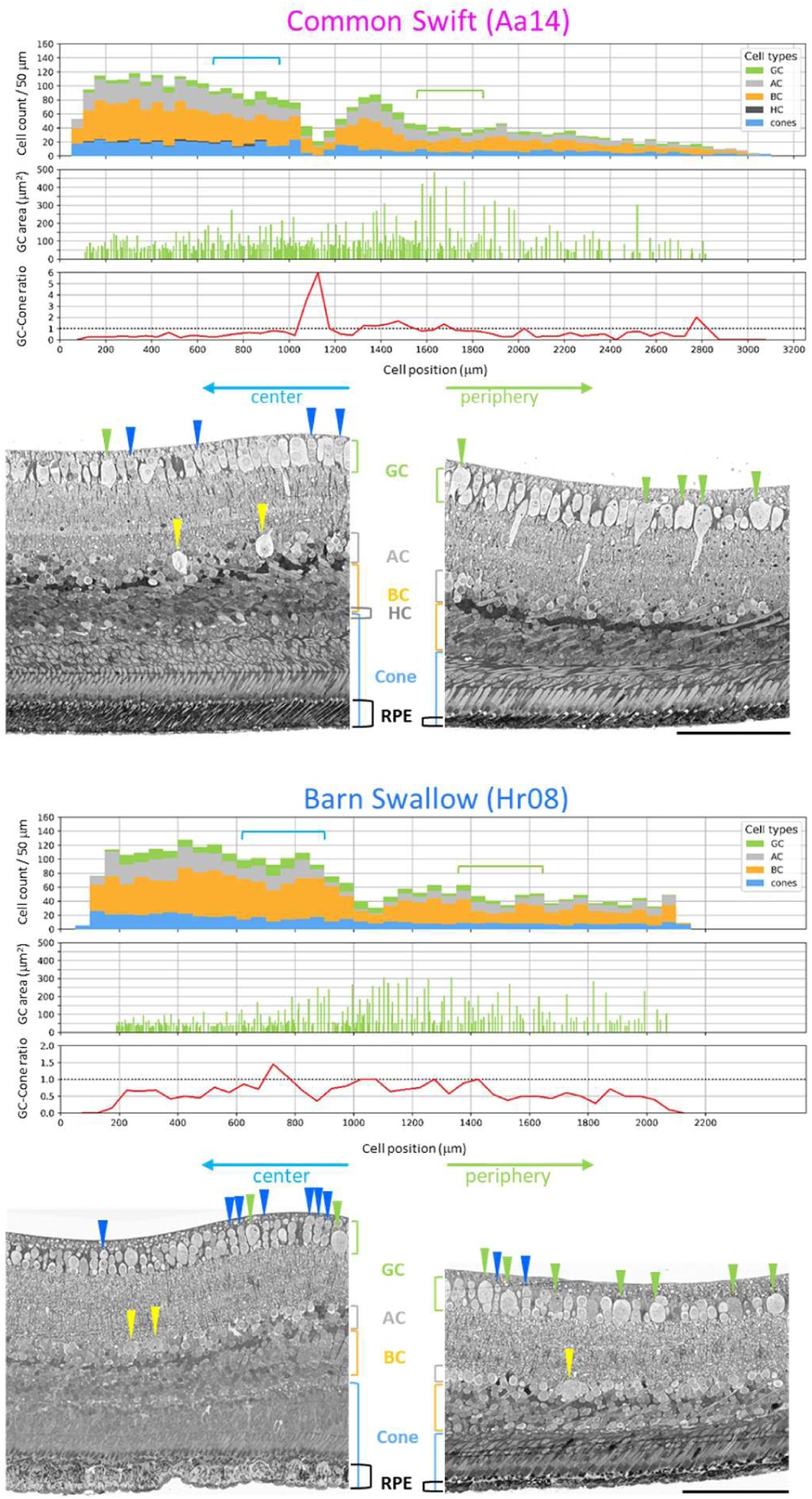
The Common Swift and Barn Swallow temporal foveae are analogous. Cone, BC, AC densities are higher in the foveal area oriented towards the center than in the area facing the periphery. Big GCs (≥ 200 μm^2^; indicated by green arrowhead) are primarily located in the area oriented towards the periphery. Numerous small GCs populate the area facing the center. These often form radial columns (blue arrowhead), though this organization is less frequent than in the central fovea (Figure 3B). Occasional large somas (yellow arrowhead) are present in the AC layer of the INL; these may represent displaced ganglion cells (DGCs). Stacked bar charts show cell counts for the Common Swift and Barn Swallow with an offset of < 5 µm and ∼50 µm, respectively, relative to the pit floor. Lateral displacement of cone nuclei results in a peak GC to cone ratio of 6:1 at the foveola. The ratio decreases to 1:1 at the foveal edge and reaches values around 0.5:1 at further eccentricity. Cell positions are measured from the section edge closest to the center. Green and blue brackets indicate the positions of the images shown on the right and left, respectively. These results were from the specimens Aa14 and Hr08. Similar results were obtained with Aa06, Hr11, and Hr21. Bar scale: 100 μm.

**FIGURE 5.**
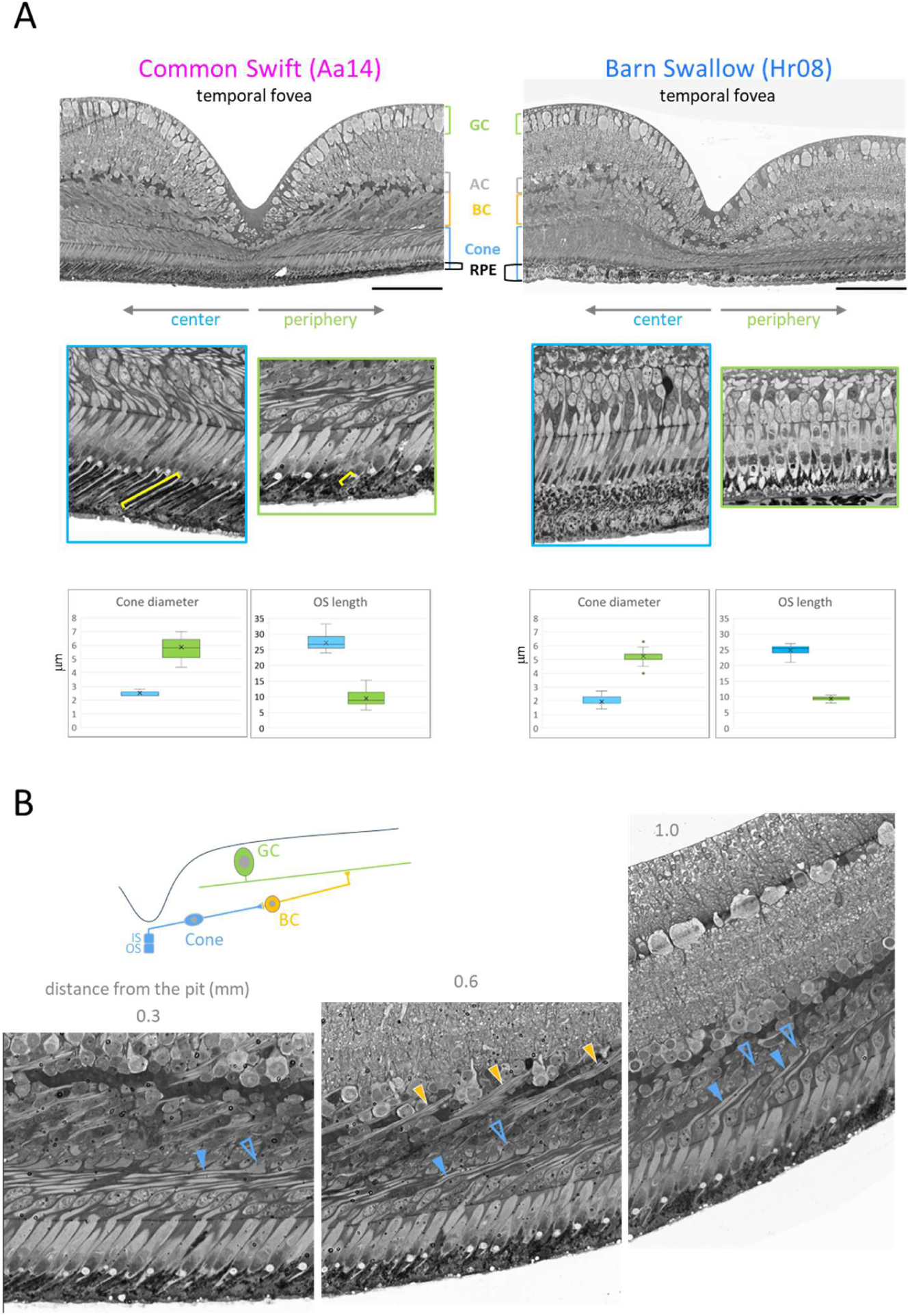
Cone sizes and axon trajectories in the temporal foveae. (**A**, upper panels) In the Common Swift and Barn Swallow, deep temporal pits arise as cone nuclei, BCs, HC, and GCs migrate laterally, leaving only the inner (IS) and outer segments (OS) embedded in the retinal pigmented epithelium (RPE) within the pit. Both RPE thickness and OS length undergo substantial changes at the foveal pit. (**A**, close up) The yellow brackets highlight the length of the OS. (**A**, lower panel) Measurement of cone diameter (at the IS level) and OS length. (**B**) In the foveal area of the Common Swift (e.g., Aa14) facing the periphery, cone axons (indicated by filled blue arrowheads) extend tangentially away from the foveal pit. Their large synaptic buttons (unfilled blue arrowhead) establish contact with BCs. The axons of these BCs (orange arrowheads) cluster together and run obliquely, ultimately connecting with the dendrites of a big GC, as shown in the schematic. Bar scale: 100 μm.

**FIGURE S3 (related to FIGURES 4 and 5).**
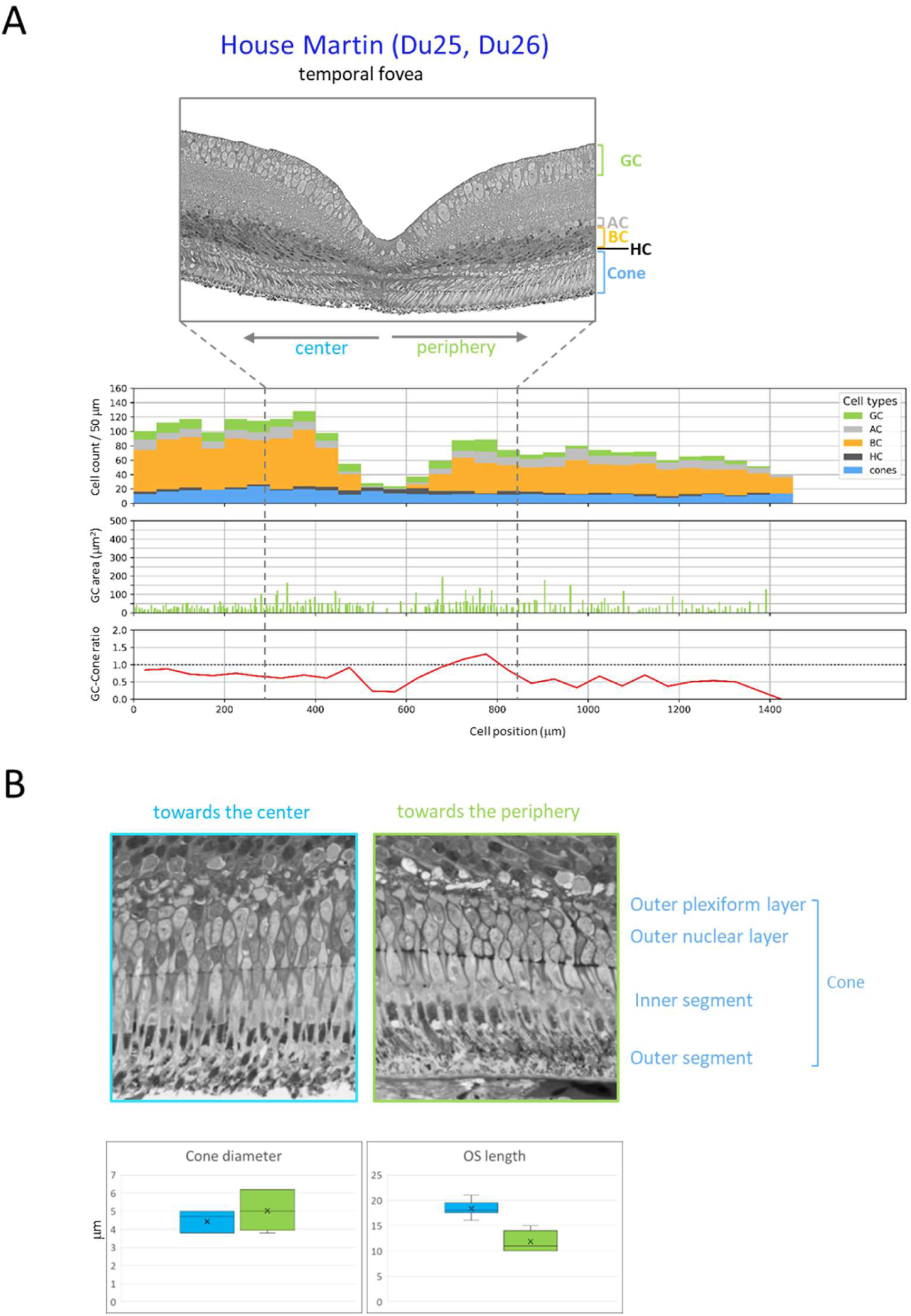
Temporal fovea anatomy in the House Martin. (**A**) Specimens Du25 and Du26 display a deep foveal pit with lateral displacement of GC, AC, BC, and cone nuclei, similar to Barn Swallows and Common Swifts, but with reduced asymmetry between the foveal area oriented towards the center and the one facing the periphery and fewer big GCs. Their sizes do not exceed 150 µm². (**B**) Cone outer segment (OS) length and inner segment (IS) diameter (thickness) measurements show less pronounced size variations between center-facing and periphery-facing cones compared to the Barn Swallow and Common Swift.

### 5. Why do raptors develop temporal foveae with shallow pits or no pit at all?

Few bird species have temporal foveae and raptors are often cited as examples. Some species of raptors have a shallow temporal fovea, while some others do not have any (Mitkus et al., 2017; Oehme, 1964; Potier et al., 2017; Rochon-Duvigneaud, 1943; Rodrigues et al., 2023). This raises two questions: what is the purpose of either a deep or shallow fovea and why does it form? To address this question we analyzed which structural elements distinguish the shallow temporal fovea of the Common Kestrel (*Falco tinnunculus*) from the dome-shaped fovea of the Black Kite (*Milvus migrans*) (Figure S4). In the Common Kestrel, the fovea had a well-formed ridge around a shallow pit. The nuclei of the foveal cones stayed in the pit while the majority of BCs and GCs moved in the parafoveal area. Cone density in the Common Kestrel’s foveal pit was ∼3 times higher than in the Black Kite’s dome-shaped fovea. In the Common Kestrel, the accumulation of cones, BCs, and GCs could result in the lateral displacement of foveal cells, creating a distinct indentation and potentially enhancing structural stability. In the Black Kite, the fovea remained dome-shaped most likely because of lower neuron densities (Figure S4). The exceptionally high densities of foveal cones and GCs in the frontal visual field should enable the Common Kestrel hovering 10–20 meters above the ground to spot prey. Notably, GCs ≥ 200 μm² were absent in the Common Kestrel, with only a few exceeding 100 μm². While present in the Black Kite, GCs ≥ 200 μm² were less numerous and more dispersed than those observed in the swallows and swifts (Figure S4). In primates, sharp frontal vison is thought to require the elongation of cone axons that make connection with BCs pushed to the side, so that the light-sensitive part of the cones gets a direct shot at the light (Masland, 2017). This notion is countered by the Common Kestrel, which effectively locates prey in its frontal visual field despite lacking a deep foveal pit and possessing a thick outer nuclear layer. Even though the lateral cell displacements observed in the fovea of primates, as well as in the temporal foveae of the Barn Swallow, the House Martin and the Common Swift could marginally increase visual acuity, we suggest that the underlying reason might be different. In swifts and swallows, the nuclei of foveal cones were displaced out of the pit and the long axons of foveal cones established connections with BCs that themselves have moved far away from the pit (Figure 5B). We suppose that these cell movements are required for enabling the wide dendritic trees of GCs ≥ 200 μm^2^, which may extend well beyond the pit, to receive input from multiple cones via laterally displaced BCs. This process does not take place in raptors because they do not have clustered big foveal GCs. However, this could occur in primates since 5-10% of foveal GCs are non-midget GC types with, for some of them, wide dendritic fields that receive input from multiple cones (Calkins and Sterling, 2007; Grunert et al., 1993; Patterson et al., 2022; Sinha et al., 2017).

**FIGURE S4 (related to FIGURES 4 and 5).**
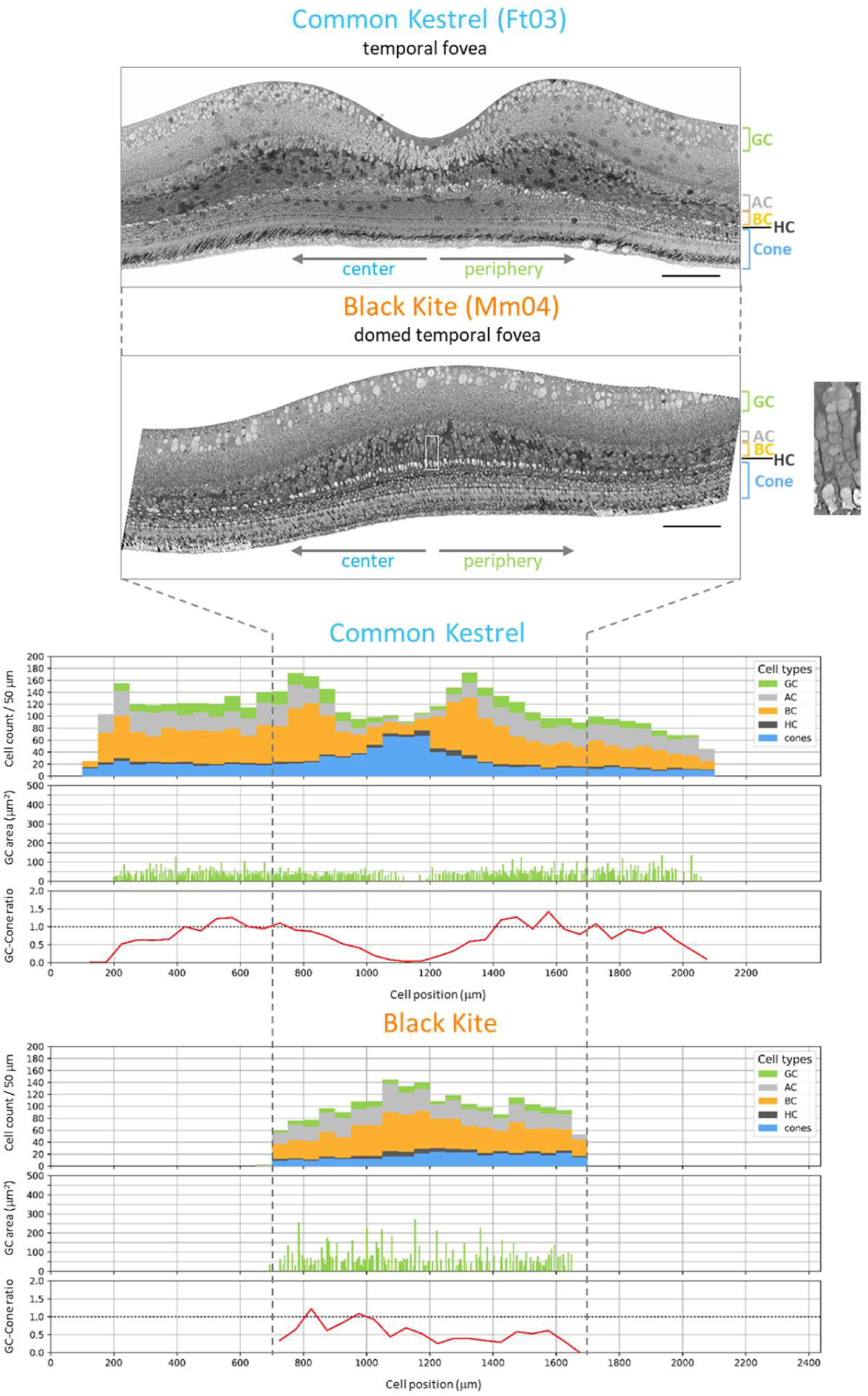
Morphological comparison of the temporal foveae in two birds of prey. In the adult Common Kestrel (Ft03), the temporal retina features a shallow foveal pit where most GCs, BCs and ACs are displaced laterally into the parafoveal area, while cone nuclei remain within the pit. In the adult Black Kite (Mm04), the fovea is dome-shaped, characterized by fewer GCs and cones compared to the Common Kestrel. The dome structure primarily results from an increase in BCs organized into distinct radial columns (close-up). Notably, while the Common Kestrel lacks GCs ≥ 200 μm² across the foveal area, the Black Kite exhibits scattered GCs of this size class (see also Table S2). Scale bars: 100 μm.

### 6. Development of a deep fovea is a slow process that involves ganglion cell apoptosis

In primates, deep fovea development is a lengthy postnatal process, taking months. In the juvenile Common Swift Aa13, the temporal foveal pit was not yet fully formed one month after this specimen had left the nest (Figure 6A, Table S1). There were clumps of GCs in place of the pit, suggesting that GCs filled the space left empty by the lateral displacement of the foveal BCs, ACs and cone nuclei. Some GCs at this location were apoptotic (Figure 6B), suggesting that the pit would have formed after the elimination of dying neurons. In older specimens (based on eye and brain sizes, Table S1), only a few supernumerary GCs remain at the edge of the pit (Aa14), and these disappear entirely by the adult stage (Aa06). In young pigeons, ∼50% of foveal GCs are eliminated at the time of the foveal pit formation (Rodrigues et al., 2023). In swifts, eliminated GCs might include those that had initially established one-to-one connections with single foveal cones via foveal BCs. The drawing by Rochon-Duvigneaud (1943; *fig. 383*) of a nestling Common Swift’s temporal retina shows exclusively small GCs, capturing the onset of lateral BC displacement alongside a slightly deepened foveal pit. Cones, BCs and ACs could be progressively recruited by GCs ≥ 200 mm^2^ during fovea maturation (Figure 6B, schema). Indeed, as detailed later in the Results section, half of the cones in the temporal fovea would eventually connect with GCs ≥ 200 μm^2^, at a 20:1 ratio. Making connections with the broad dendritic fields of these big foveal GCs would require foveal BCs to move laterally and cone nuclei to develop long axons (Figure 5B).

**FIGURE 6.**
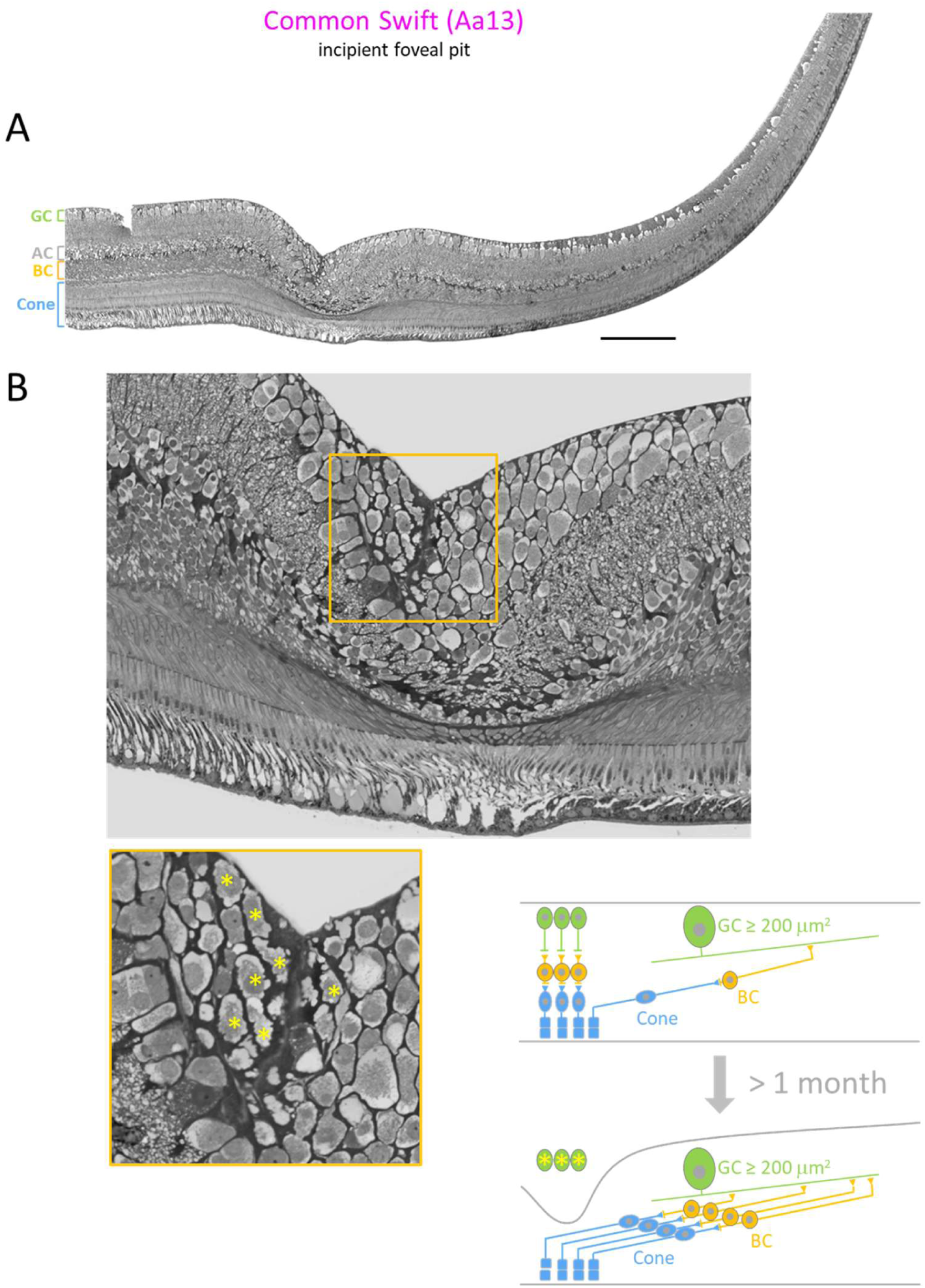
Temporal fovea maturation. (**A**) Temporal fovea of the Common Swift Aa13. This specimen lived at least one month after leaving the nest (Table S1). Cone nuclei, BCs and ACs have completed their lateral movement. Despite this, a fully formed pit does not develop as GCs occupy the space left by cell displacement. (**B**) Close-up of the incipient pit: The somas filling the pit (marked with yellow asterisks) exhibit highly irregular membranes with bulging protrusions; such membrane blebbing is a hallmark of apoptosis. Schema: During the month-long maturation of the fovea, developmental refinement occurs. Cones and BCs initially synapse with small GCs before reorganizing. They shift laterally away from the foveal center to connect with bigger GCs characterized by broad dendritic fields. This remodeling is driven by the apoptosis of the original small GCs and an increased number of big GCs. Scale bars: 200 μm.

### 7. How does the temporal foveae compare with non-foveate temporal retinas?

Apart from the pit, one might wonder if the swift and swallow temporal foveae possess other characteristics that distinguish them from non-foveate temporal retinas. The Eurasian Sparrowhawk (*Accipiter nisus*), the Grey Wagtail (*Motacilla cinerea)*, and the Eurasian Blackcap (*Sylvia atricapilla*) are birds with a central fovea but with no temporal fovea (Figure S5A). The hawk and the two passerines differ in many aspects, but, despite this, their temporal retinas were similar. Cones were uniformly thick with short OS (Figure S5B) resembling cones found in the swift and swallow temporal foveal areas facing the periphery (Figure 5A). The three species displayed high densities of cells in the GCL and the ratios of cones to GCL’s cells fluctuated around 1:1. Somas ≥ 200 mm^2^ were rare in the GCL of the two passerines and absent in the hawk (Figure S5A). The convergence ratios between BCs and GCs were similar in the non-foveate temporal retinas and in the foveal areas facing the periphery, i.e., 3.7 in An01, 4.5 in Mc12, 4.5 in Sa09, 3.7 in Aa14, 4.5 in Hr08. These values were significantly lower than those in the swift and swallow central retinas, i.e., 24 in Aa14, 10 in Hr08 (Figures 2B, 3B). Taken together, our results suggest that a circuit structure where 1 cone connects through 4 BCs to 1 GC could be one of the main circuit organization in the non-foveate temporal retinas of these birds. One might wonder why the two passerines do not develop a temporal fovea, whereas the House Martin (also a passerine) develops one (Figure S3). One reason could be that the ratio of GCs to cones was 9:10 in the temporal retinas of the Eurasian Blackcap and Grey Wagtail, whereas it was 6:10 in the House Martin’s temporal fovea. This disparity is significant enough to suggest that, unlike the House Martin, the Eurasian Blackcap and Grey Wagtail possess fewer GCs that receive input from multiple cones via laterally displaced BCs, thus preventing the development of a foveal pit.

**FIGURE S5 (related to FIGURES 4, 5 and 6).**
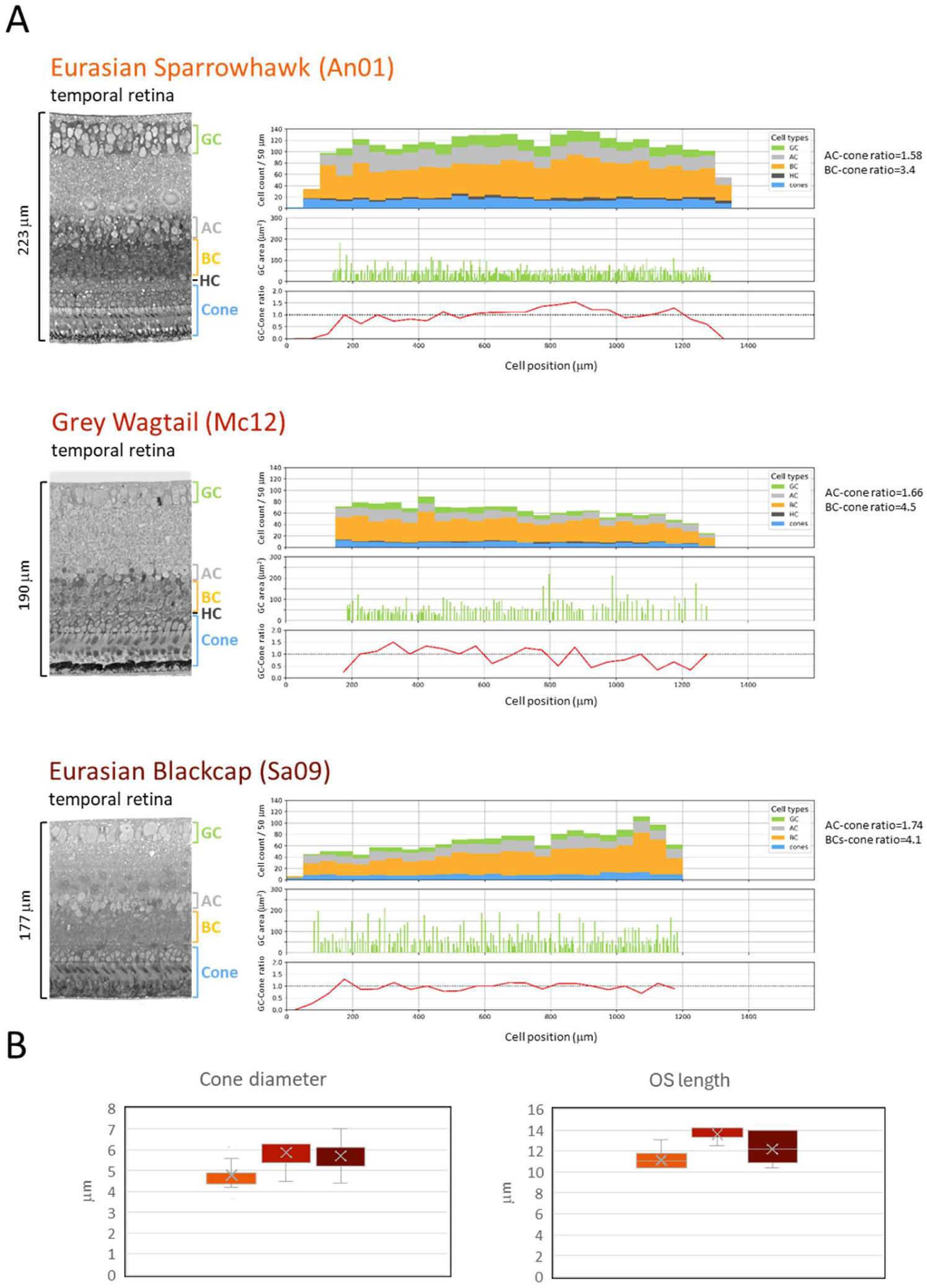
Organization and photoreceptor morphology of non-foveate temporal retinas in birds with a central fovea. (**A**) Comparative retinal organization across three species from distinct taxonomic Family and Order. Note the higher peak densities of cones, BCs, ACs, and GCs in the Eurasian Sparrowhawk. Across all three species, the GC-to-cone ratio remains consistently near 1:1 along the entire retinal fragment. Big GCs (≥ 200 µm²) are absent in the hawk and rare in the passerines. (**B**) Morphometric analysis of cone inner segment (IS) thickness and outer segment (OS) length.

### 8. Clusters of big ganglion cells, a specific feature of the temporal foveae

Measurements in the central, temporal and peripheral retinas of 8 species led us to conclude that the mean areas of somas in the GCL ranged from 35 μm^2^ to 55 μm^2^ and that somas ≥ 200 μm^2^ were rare (Figure S6; Table S2). It was only in the temporal foveae of the Common Swift and Barn Swallow that clusters of GCs ≥ 200 μm^2^ were identified. They accumulated in the vicinity of the foveal pit, and more specifically in the area oriented towards the periphery (Figure 7). This area is 0.7 mm^2^ in the Barn Swallow and 1.0 mm^2^ in the Common Swift and represents 1-2% of the total retinal surface. In these foveal areas, the ratios of GCs to cones were around 1:2 in swift and swallow (Figure 7), whereas they were around 1:1 in non-foveate temporal retinas (Figure S5A). Lower ratios in the fovea could result from the loss of GCs during formation of the foveal pit (Figure 6), suggesting that the development of a population of big foveal GCs might correlate with a decrease in the number of smaller GCs. In the foveal area of the Barn Swallow, there were 417 GCs ≥ 200 μm^2^ out of a total of 10’900 GCs and 18’700 cones, and in the Common Swift, there were 673 GCs ≥ 200 μm^2^ out of a total of 8’500 GCs and 22’000 cones (Figure 7). This suggests that most of the foveal GCs were connected to cones in a ratio of 1:1 and that the number of cones available for the GCs ≥ 200 μm^2^ might not exceed 8 × 10^3^ in the Barn Swallow and 13 × 10^3^ in the Common Swift, i.e., ∼20 cones on average per GC ≥ 200 μm^2^. However, this number might be an overestimate given that we did not take into account here GCs with sizes between 100 μm^2^ and 200 μm^2^, which were fairly numerous (Figures 4, S6) and could receive input from several cones. We are faced with a paradoxical situation where GCs ≥ 200 μm^2^ and their putative broad dendritic arbors clustered in a retinal area that displayed the highest GC to cone ratios. Cajal et al. (1972; plate VI, Fig. 16) identified an orthotopic parasol-like GC connected to five cones via five BCs in the fovea centralis of an European Green Finch (*Chloris chloris*). More recently, it has been suggested that there is no avian equivalent of parasol cells as described in mammals (Seifert et al., 2023; Yamagata et al., 2021), a conclusion based on data obtained in the non-foveate Domestic Chicken. In the absence of information on the molecular properties of these big GCs in swifts and swallows, we name them *foveal magnocellular-like ganglion cells (FMGCs)*.

**FIGURE 7.**
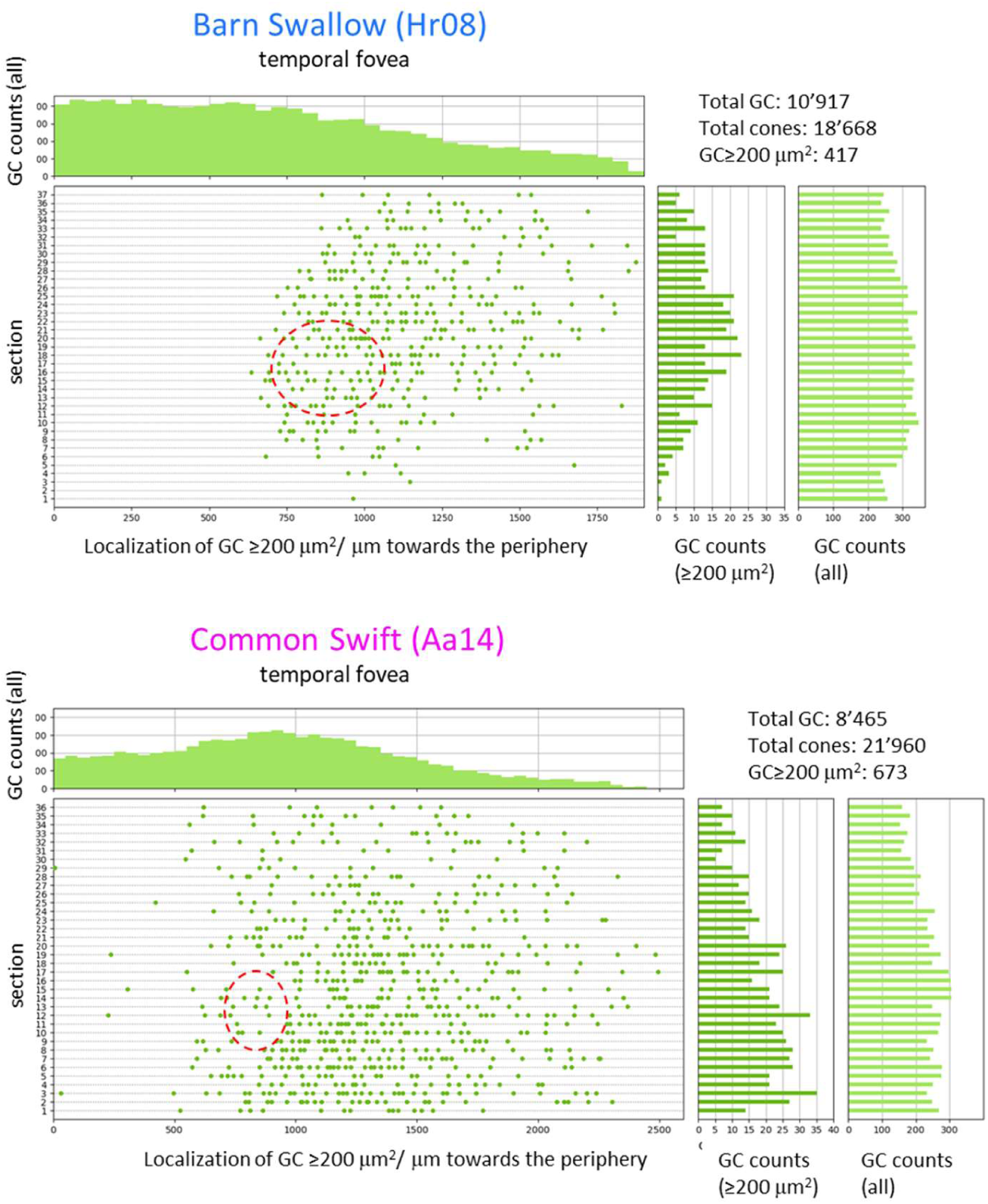
Mapping big ganglion cells (FMGC) in the temporal foveae of the Common Swift and Barn Swallow. GCs were identified across 36–37 serial semi-thin sections (1 µm) of the temporal fovea, sampled at 20–25 µm intervals. Each green dot indicates a GC with a soma area ≥ 200 µm². The adjacent histograms show the count of these big GCs alongside the total number of cell bodies within the GCL for each corresponding section. Upper-panel histograms show the total GCL cell counts across 36 or 37 sections in 50 µm sectors. No cell bodies ≥ 200 µm² were detected in the INL within these sections. Due to their small size (5–7 µm diameter), cone nuclei could not be fully quantified across the serial sections. The total number was estimated by multiplying the cone density per unit area (section length [µm] × 1 µm) by the width of the foveal region. Average GC to cone ratios across the entire foveal surface—0.6 in the Barn Swallow and 0.4 in the Common Swift—mirror those found in individual sections. This consistency indicates that the vast majority of foveal GCs were captured in the serial sections. Red circles mark the positions of the foveal pits.

**FIGURE S6 (related to FIGURE 7).**
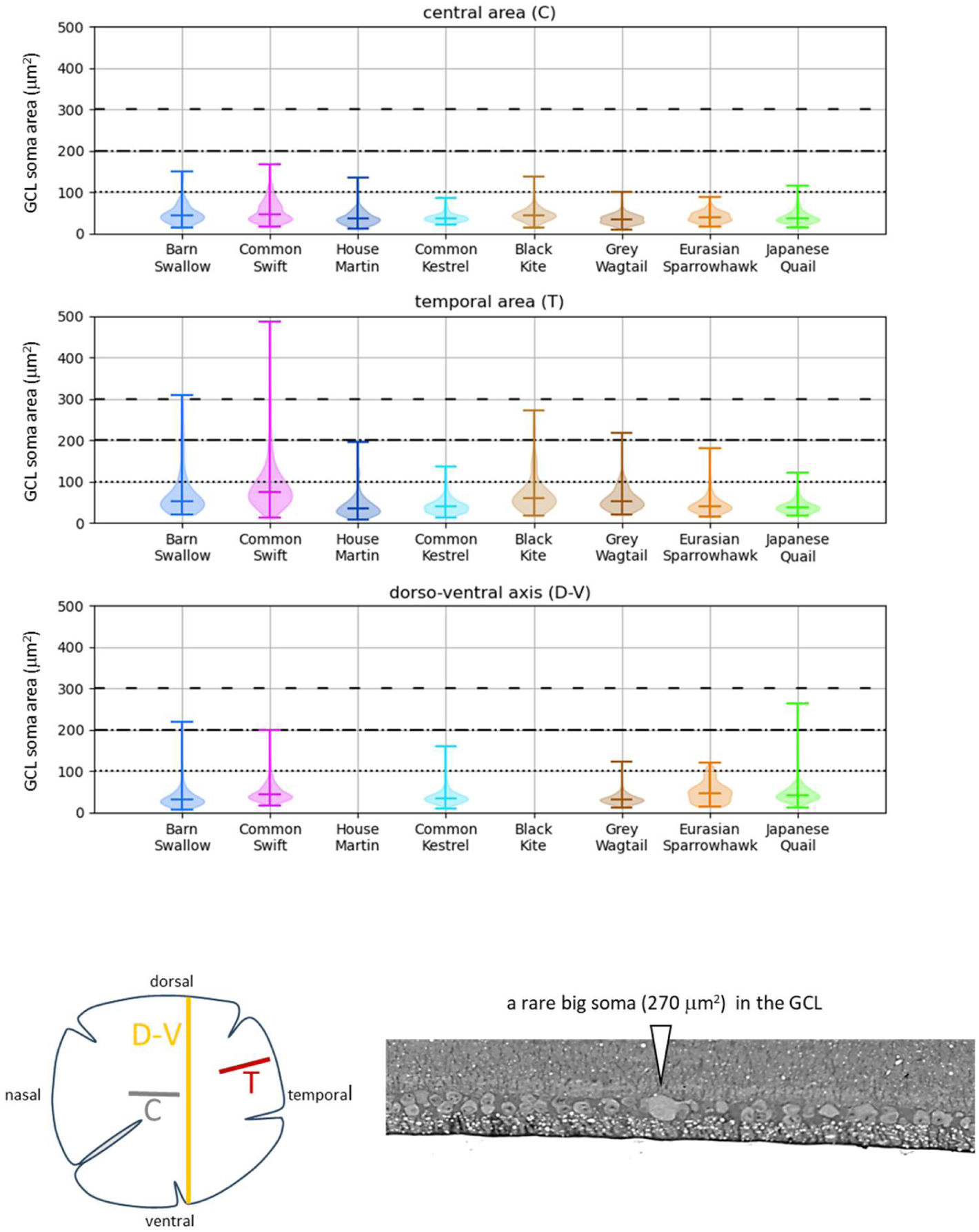
Spatial distribution and morphological characteristics of GCL somata in the avian retina. GCs with a soma area ≥ 200 μm² are generally rare and solitary across the species examined. Extensive analysis of the ganglion cell layer (GCL) reveals an absence of these big cells in the central retina (C). Along the dorsoventral (D-V) axis, only 4 GCs ≥ 200 μm² were identified among 5,426 total cells spanning 49 mm across six species. Notably, clusters of big GCs are found exclusively within the temporal areas (T) of the Barn Swallow and the Common Swift. Micrograph showing a unique large soma (270 µm²) within a population of 806 GCL cells spanning 10 mm across sections C, T, and D-V in the Japanese Quail. See also Table S2.

**TABLE S2 (related to FIGURE 7).** Cell number and cell size in the Ganglion Cell Layer.

|  |  | GCL, total<br>number of cells | GCL length<br>(μm) | Median cell area<br>(μm<sup>2</sup>) | Cell area |  |  |
| --- | --- | --- | --- | --- | --- | --- | --- |
|  |  |  |  |  | ≥100 μm<sup>2</sup><br>(%) | ≥200 μm<sup>2</sup><br>(%) | ≥300 μm<sup>2</sup><br>(%) |
| Barn Swallow | C | 267 | 774 | 46 | 6 | 0 |  |
|  | T | 157 | 731 | 71 | 29 | 10 | 5 |
|  | D-V | 929 | 8'293 | 31 | 2 | 0.2 | 0 |
| Common Swift | C | 302 | 3'517 | 46 | 8 | 0 |  |
|  | T | 315 | 2'987 | 75 | 30 | 6 | 3 |
|  | D-V | 578 | 8'106 | 44 | 4 | 0.2 | 0 |
| House Martin | C | 427 | 1'743 | 35 | 1 | 0 |  |
|  | T | 261 | 1'434 | 35 | 5 | 0 |  |
|  | D-V | N/A |  |  |  |  |  |
| Common Kestrel | C | 168 | 688 | 37 | 0 |  |  |
|  | T | 512 | 1'900 | 40 | 2 | 0 |  |
|  | D-V | 1'372 | 12'000 | 36 | 1 | 0 |  |
| Black Kite | C | 462 | 1'340 | 44 | 1 | 0 |  |
|  | T | 169 | 955 | 60 | 15 | 3 | 0 |
|  | D-V | N/A |  |  |  |  |  |
| Grey Wagtail | C | 260 | 621 | 34 | 0 |  |  |
|  | T | 159 | 1'109 | 52 | 8 | 1 | 0 |
|  | D-V | 1'600 | 7'800 | 32 | 0 |  |  |
| Eurasian Sparrowhawk | C | 169 | 703 | 40 | 0 |  |  |
|  | T | 370 | 1'108 | 40 | 2 | 0 |  |
|  | D-V | 141 | 2'061 | 47 | 6 | 0 |  |
| Japanese Quail | C | 331 | 2'417 | 36 | 0 |  |  |
|  | T | 155 | 1'240 | 38 | 1 | 0 |  |
|  | D-V | 806 | 10'800 | 43 | 3 | 0.1 | 0 |

### 9. An algorithm for analyzing the topographic distribution of optic nerve axons

The study of ganglion cell axons, specifically those originating from the fovea, is essential for understanding how visual information is transmitted from the retina to the brain. We first wanted to determine axon numbers and distribution in the ON. Studies on bird’s ON are sparse and concern species which can be counted on the fingers of one hand (Binggeli and Paule, 1969; Ikushima et al., 1986; O’Flaherty, 1971; Rager, 1980; Wathey and Pettigrew, 1989). To comprehensively analyze the topographic distribution of axons, investigating the entire nerve surface is necessary to map axon counts and axon sizes. Analyzing one ON of one Barn Swallow specimen with TEM would require > 2 × 10^3^ images like the one presented in Figure S7. In most studies, axon densities and axon counts are estimated from small random samples. We chose an approach where high quality semi-thin sections of the transversely cut ONs were analyzed in their entirety by light microscopy. In one image of Barn Swallow’s ON (Figure 8A), there are 2.35 × 10^5^ axons ≥ 0.1 μm^2^ and manually surround each one using an image processing program would have required > 235 hours. To reduce analysis time to a few hours, we have developed and trained an algorithm to create maps of axon densities and axon sizes (Figure 8B). The algorithm can detect axons > 0.1 μm^2^ (0.4 μm in diameter). TEM analysis indicated that axons < 0.1 μm^2^ represent 12% of all axons (Figure S7). The performance of the algorithm was tested for each ON analyzed. On average, it detected 86.6% of axons with sizes between 0.1 μm^2^ and 0.5 μm^2^ and 92.9% of axons > 0.5 μm^2^ (Figure S8).

**FIGURE 8.**
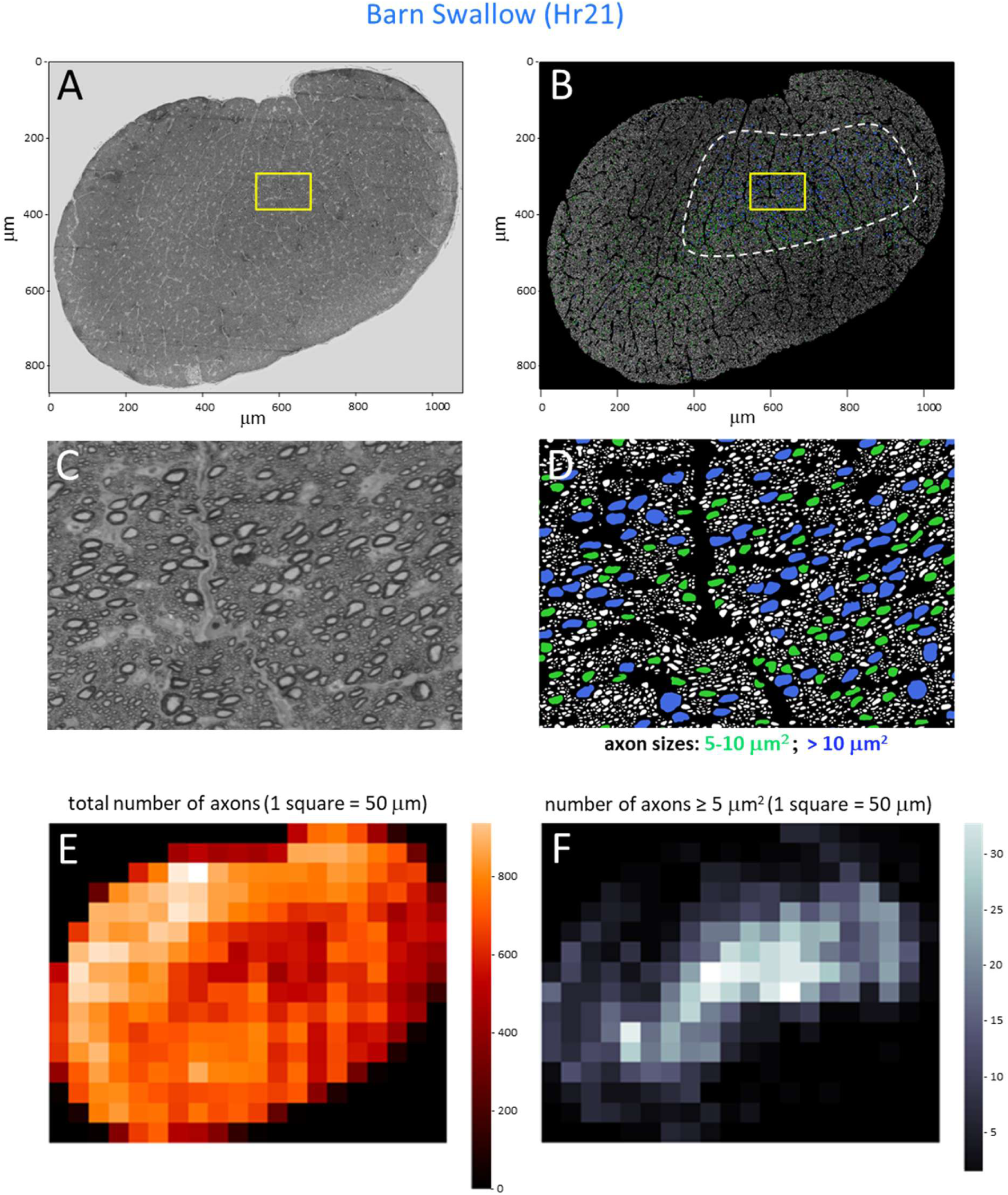
Axon distribution and axon size in the ON of the Barn Swallow. (**A**) Optical microscopy image of a cross-section of the ON. (**B**) algorithm output. Axons are color-coded by size: green (5–10 µm²), blue (> 10 µm²), and white (< 5 µm²). While the green and blue distributions partially overlap, the blue axons occupy a smaller overall area; specifically, over 90% of them are clustered within a region bounded by the closed dashed line, which represents 22% of the total cross-sectional area. Yellow rectangles in **A** and **B** delimit the magnified views shown in **C** and **D**, respectively. (**E**, **F**) Heatmaps of the optic nerve generated by the algorithm to visualize axon density (**E**) and axon size (**F**) distributions.

**FIGURE S7 (related to FIGURE 8).**
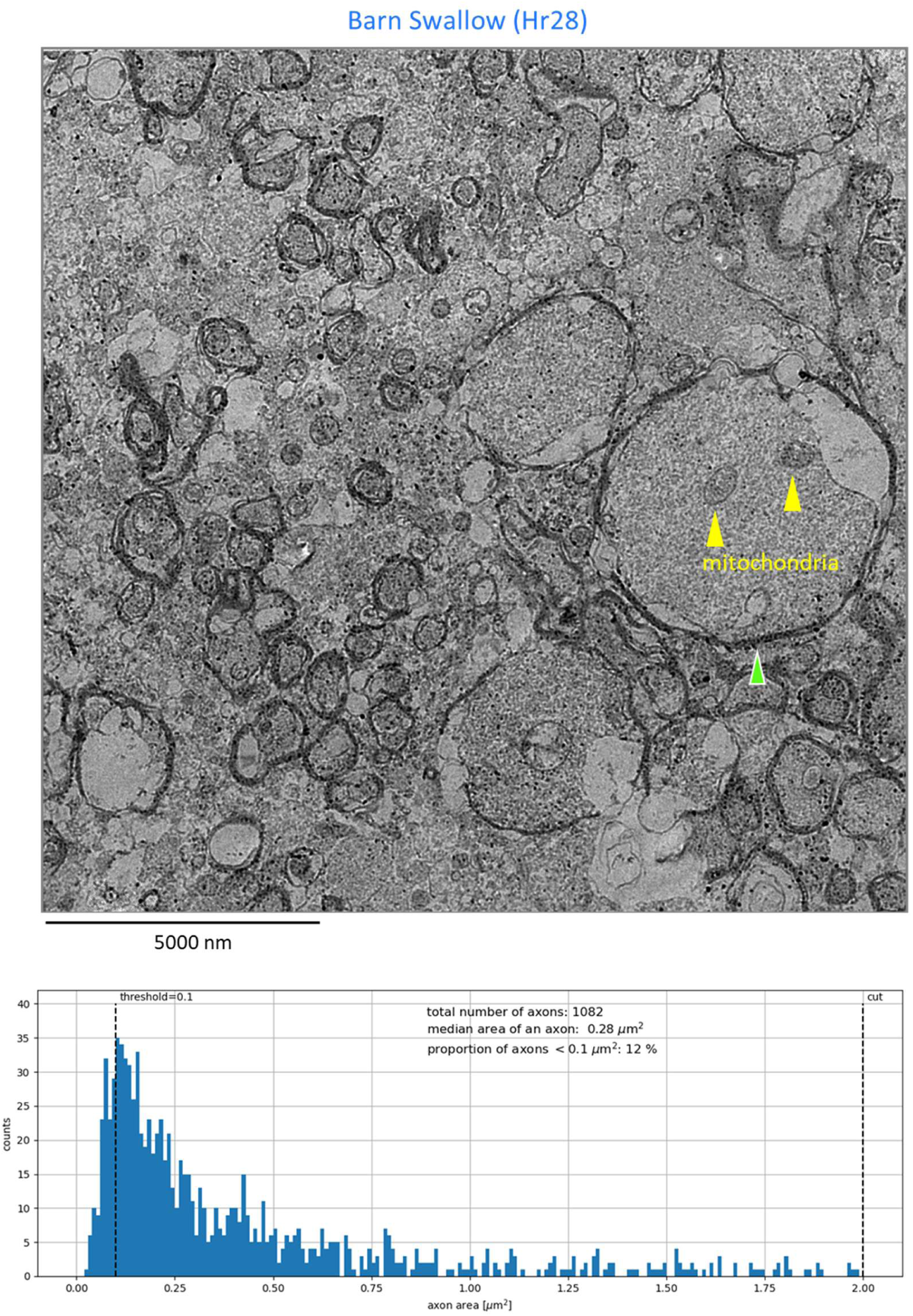
Ultrastructural analysis of the adult Barn Swallow optic nerve (ON). Transmission electron microscopy (TEM) of a central sector of the ON. Myelinated axons were manually outlined for morphometric analysis. Of the 1,082 myelinated axons measured, 12% possess a surface area < 0.1 μm², which represents the detection threshold for semi-thin sections analyzed via light microscopy. The green arrowhead indicates an axon of 20 μm² (5 μm in diameter), representing one of the largest axon calibers observed in the avian ON.

**FIGURE S8 (related to FIGURES 8, 9 and 10).**
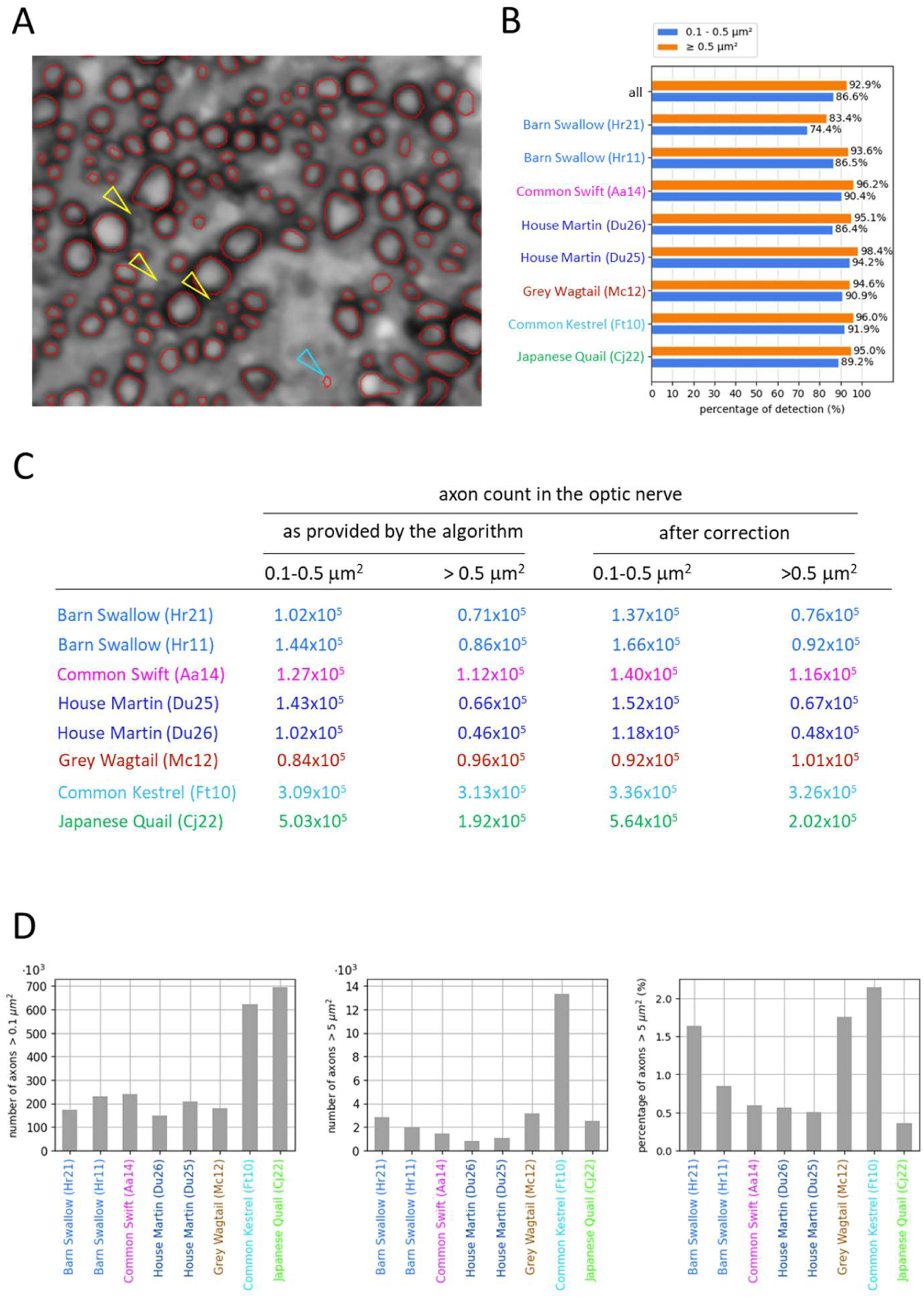
Performance assessment of the axon segmentation algorithm. (**A**) Representative images used for manual validation. For each ON, 15 randomly selected images were manually edited. While the algorithm accurately identifies most axons (red circles), errors include the omission of small-caliber axons (yellow arrowheads) and occasional false positives (blue arrowheads). (**B**) Proportions of axons correctly identified by size category (0.1–0.5 µm² and > 0.5 µm²). The decrease in algorithmic performance is primarily restricted to axons < 0.5 µm² due to blurred morphological outlines. (**C**) Correction of total axon counts (≥ 0.1 µm²) per optic nerve. Automated counts were adjusted by adding the estimated number of missed axons identified in (**A**) and (**B**). After accounting for 12% of small, undetectable axons (< 0.1 µm²) (Figure S7), total axon counts reached 2.35 × 10⁵ for the Barn Swallow (Hr21) and 2.78 × 10⁵ for the Common Swift (Aa14). Given that these species possess fewer than 3’000 centrifugal fibers (Feyerabend et al., 1994), we conclude that 98% of these axons originate from GCs. (**D**) Interspecies comparison of axon size distributions.

### 10. Large axons cluster together in the optic nerves of swallows and swifts

We have established maps of axon densities and axon sizes (Figures 8, 9, 10B, S9A) and quantified size distribution in the ONs of seven bird species (Figures 10A, S9B). We wondered whether axons that originate from GC ≥ 200 μm^2^ (FMGC) localized in the swift and swallow temporal foveae could be identified and displayed specific physical arrangements. In the Barn Swallow specimen Hr21, there were 417 GCs ≥ 200 μm^2^ in the temporal fovea (Figure 7) and 487 axons ≥ 10 μm^2^ were counted in the ON (Figures 8, 10B). The regions occupied by mid-sized (5–10 µm²) and large (> 10 µm²) axons partially overlapped. Notably, 91% of the large axons (444 count) clustered within a well-defined area representing 22% of the total ON area. In the Common Swift specimen Aa14, there were 673 GCs ≥ 200 μm^2^ in the temporal fovea (Figure 7). Among the 69 axons > 10 μm^2^ counted in the ON, 64 were localized in an area corresponding to 9% of the total ON area (Figure 10B). 594 mid-sized axons (5-10 μm^2^) were localized in this delimited area, accounting for half of the mid-sized axons in the ON. Therefore, 658 axons originating from the temporal fovea were grouped together despite the fact that they were on average smaller than those of the Barn Swallow. The number of axons > 10 μm^2^ was similar in the adult specimen Aa06 (data not shown). The Barn Swallow specimen Hr11 was expected to be younger than the specimen Hr21 (Table S1). While the number and size distribution of axons > 0.5 μm^2^ were quite similar in Hr11 and Hr21 (Figures 10A, S8C), they were more axons > 10 μm^2^ in Hr21 than in Hr11 (Figure 10A, B). In Hr11, there were 226 axons > 10 μm^2^ grouped in an area corresponding to 22% of the total ON area, much like in Hr21. In Hr11, we suppose that a fraction of axons that originated from the temporal fovea and who were regrouping with the 226 axons > 10 μm^2^ (Figure 10B) had not yet reached their adult sizes. In the ONs of the House Martin specimens Du25, Du26 (> 5 month-old, Table S1), 86% (Du25) and 91% (Du26) of axons > 10 μm^2^ were grouped in an area corresponding to 20% of the total ON area. However, there were 5 to 8 times fewer axons > 10 μm^2^ in the House Martin than in the Barn Swallow (Figures 9, 10B), in line with the fact that there were fewer big GCs in the House Martin temporal foveae and that their sizes did not exceed 150 μm^2^ (Figure S3). Noteworthy, the total numbers of axons ≥ 5 μm^2^ in the areas that regroup ∼90% of the axons > 10 μm^2^ were the same in Du25 and Du26 (Figure 10B).

**FIGURE 9.**
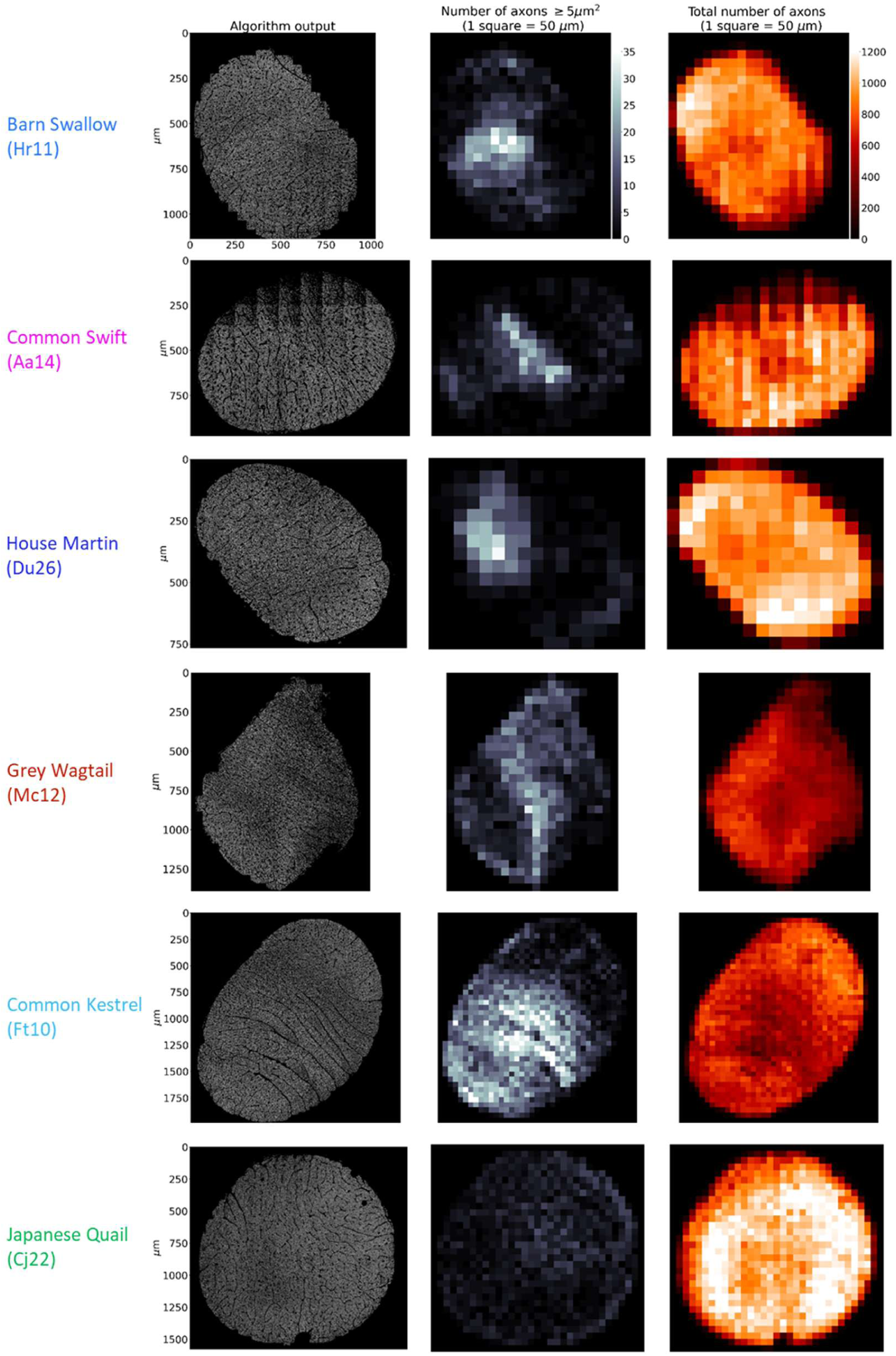
Interspecies comparisons reveal significant heterogeneity in axon size and density across the ON. The Barn Swallow, House Martin, and Common Swift differ from the Grey Wagtail and Common Kestrel by the topographic clustering of large axons (≥ 5 µm²) within circumscribed regions and their near-absence at the periphery. The Japanese Quail, a Galloanserae, is characterized by high density of small-diameter axons and a low frequency of axons ≥ 5 µm² distributed across the entire ON surface.

**FIGURE 10.**
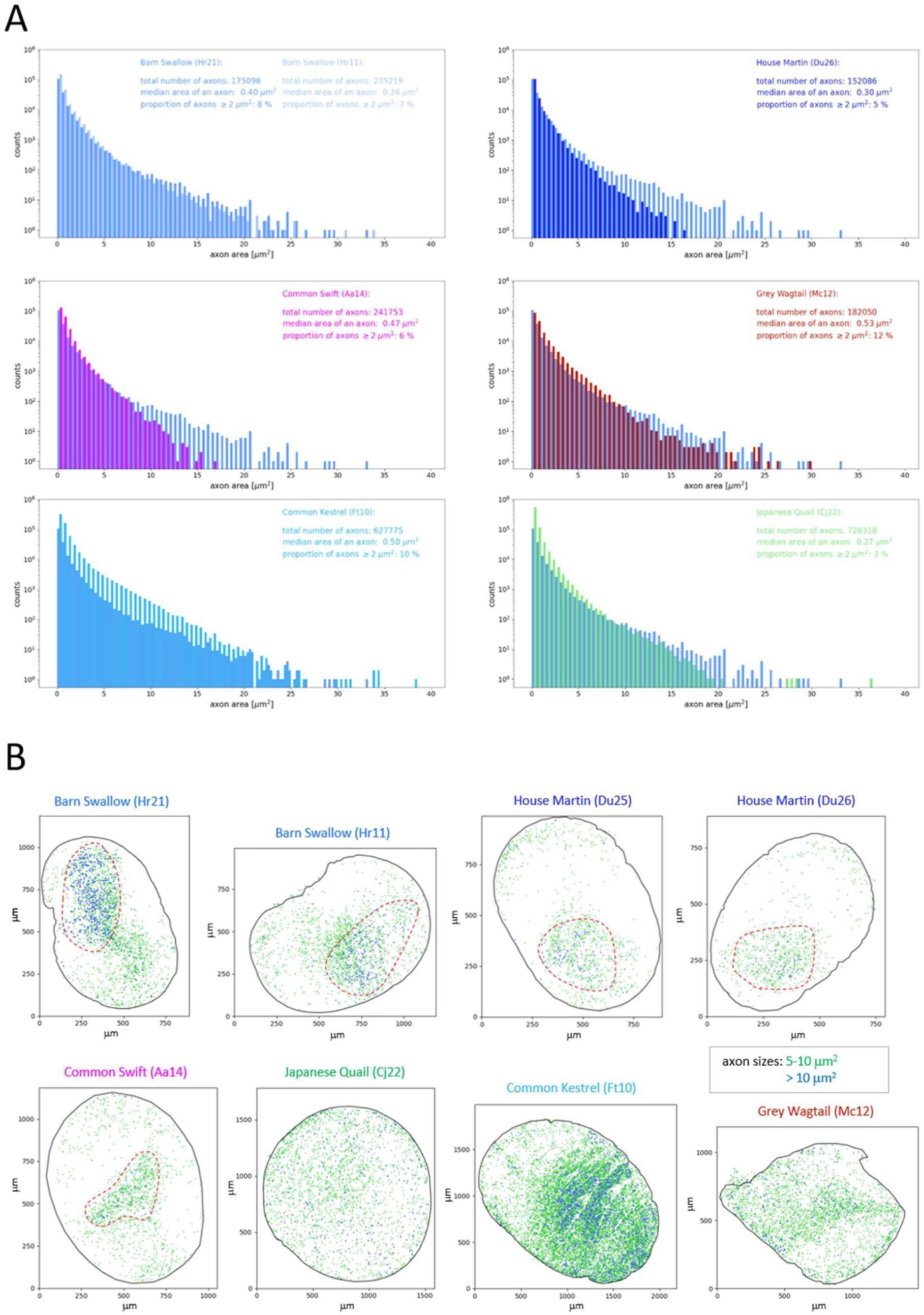
Quantified size distribution of axons (≥ 0.1 µm²) in the ON. (**A**) Size distribution as determined by the algorithm. Each specimen or species is compared to the Barn Swallow (Hr21). Corrected values for the total number of axons, adjusted for algorithm performance, are listed in Figure S8C. (**B**) Topographic distribution of mid-sized (5–10 µm², green) and large (>10 µm², blue) axons in the ON. In the Barn Swallows (Hr21, Hr11), a delimited area representing 22% of the total ON surface contains 91–92% of all blue axons. In the Common Swift (Aa14), a smaller area (9% of total) encompasses 92% of blue and 50% of green axons (658 axons ≥ 5 µm²). In each of the two House Martin specimens (Du25, Du26), an area representing 20% of the surface of the ON contains 86-91% of blue axons and totals 460 axons (blue and green). Conversely, in the Grey Wagtail and the Japanese Quail, blue axons are uniformly sparse across the entire surface of the ON, with no areas of increased density. In the Common Kestrel, dense populations of green and blue axons cover over two-thirds of the ON surface; notably, axons > 10 μm^2^ cluster into three distinct high-density zones.

### 11. Large axons do not cluster in the optic nerves of birds with no deep temporal fovea

We wondered whether the clustering of large axons in the ON was related to the presence of a temporal fovea. In the non-foveate Japanese Quail, rare axons > 10 μm^2^ were spread out throughout the entire surface of the ON and showed no tendency to cluster (Figures 9, 10B). The most interesting species to compare with the swallows and the swifts were Falconidae like the Common Kestrel and the Peregrine Falcon (*Falco peregrinus*), which possess shallow temporal foveae (Figure S4; Mitkus et al., 2017; Rodrigues et al., 2023). Their ONs had the highest number and proportion of axons > 5 μm^2^ and > 10 μm^2^ but they did not regroup in a single specific location (Figures 10B, S9A). This went with the fact that there was no GC ≥ 200 μm^2^ in the temporal fovea (Figure S4). In the Grey Wagtail, there was also high number and high proportion of axons > 5 μm^2^ but they were scattered (Figures 9, 10B). In quail, kestrel and wagtail, extensive screening of different retinal areas did not reveal clusters of big somas in the GCL. They were always solitary (Figure S6; Table S2). Taken together, our results suggest that the clustering of large axons in the ON of swallow and swift was primarily driven by their shared origin from the population of GCs ≥ 200 μm^2^ (FMGC), specifically localized in the temporal foveae.

**FIGURE S9 (related to FIGURES 9 and 10).**
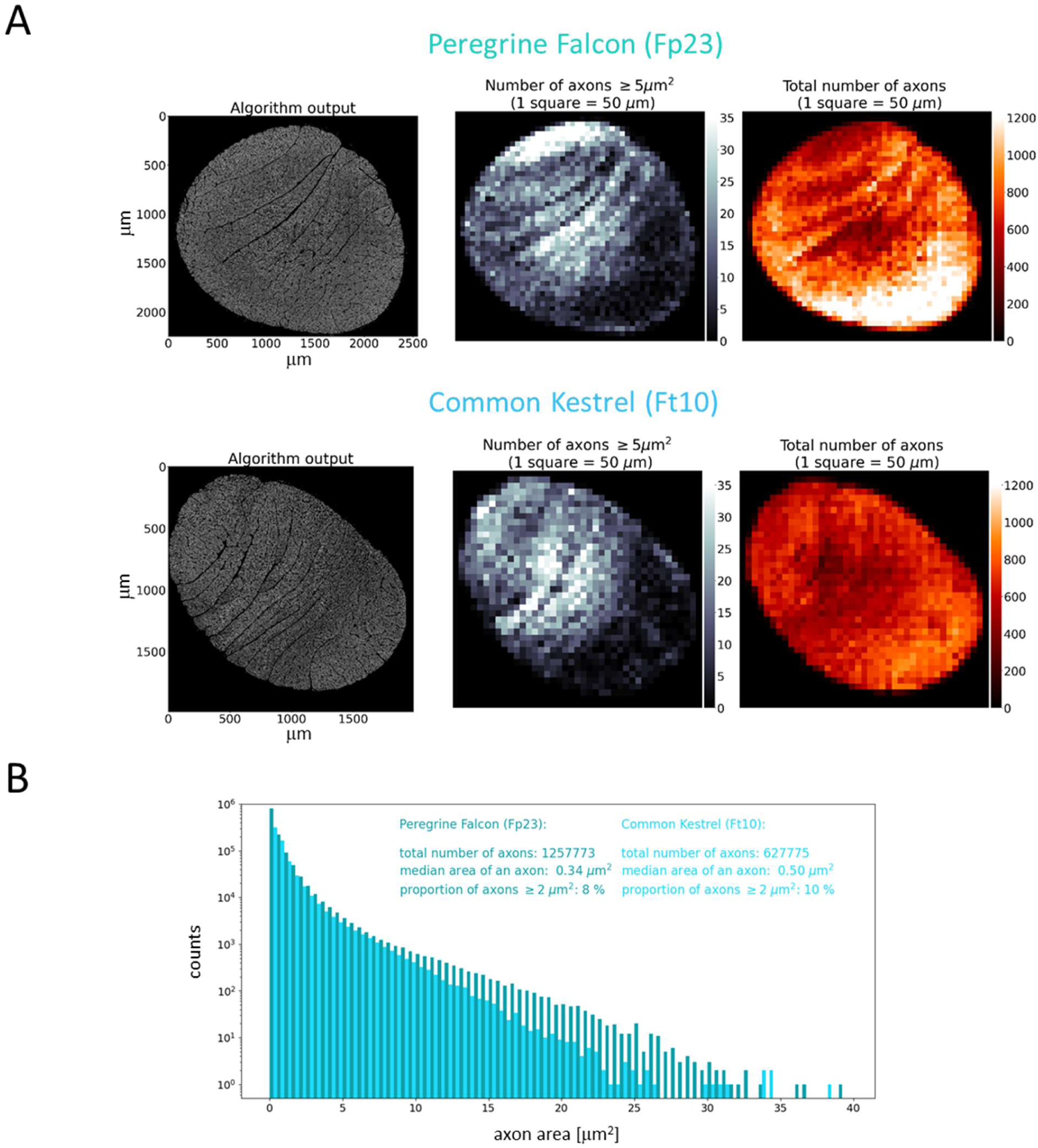
Comparative analysis of optic nerve (ON) morphology in the Peregrine Falcon and Common Kestrel. (**A**) Spatial distribution of ON axons across nerve compartments. Despite differences in species size (Table S1) and hunting behaviors (Figure 1B), both Peregrine Falcon and Common Kestrel exhibit similar distribution patterns of axon density and size. Axons with a large-sectional area (≥ 5 µm²) are distributed throughout compartments defined by connective tissue septa over two-thirds of the ON surface. (**B**) Quantitative comparison of total axon populations. The Peregrine Falcon possesses approximately twice the number of ON axons compared to the Common Kestrel, a disparity driven primarily by a greater abundance of axons with a small cross-sectional area (0.1-0.5 µm²). Total ON surface area is 3.87 mm² for Peregrine Falcon and 2.46 mm² for Common Kestrel.

### 12. The Barn Swallow’s optic nerve might be the fastest for transmitting visual information from the frontal visual field

The Common Kestrel and Japanese Quail had fairly similar number of axons in their ONs, while the proportion of axons ≥ 5 μm^2^ was about 6 times higher in the kestrel (Figures 9, S8D). This, despite the fact that the somas of the GCs in the retinas of both species were in the same size range (Figure S6). Axon diameter growth is a major driver of increases in conduction speeds along myelinated axons that requires major molecular adaptation (Bin et al., 2024). Thicker optic nerve axons could be a significant evolutionary trait in Neoaves compared to Galloanserae. The Barn Swallow, the Grey Wagtail and the Common Kestrel have the highest proportions of myelinated axons ≥ 5 μm^2^ (Figures 10A, B, S8D). However, it was only in the Barn Swallow that axons ≥ 10 μm^2^ were clustered together and originated from one retinal specialty, i.e., the temporal fovea. To date, members of Hirundinidae are unique among passerines (song birds) for possessing a temporal fovea. Passeriformes are thought to have appeared 20 million years after the Apodiformes, which includes the swifts (Brown and Mindell, 2009). It is plausible that the swallows have re-activated a pre-existing program for the development of the temporal fovea initially established by the swifts. The Barn Swallow temporal fovea is more closely analogous to that of the Common Swift than that of the House Martin, suggesting that replicating the swift’s temporal fovea, and even improving it further by developing thicker axons, should have been a complex challenge that the House Martin has not achieved with the same success. In that species, the proportion of axons ≥ 5 μm^2^ was smaller than in other species, but yet the largest axons were grouped in an area of similar size than in the Barn Swallow (Figure 10B). Even with fewer and smaller FMGCs than in the Barn Swallow, the House Martin’s foraging and flight performances are far superior to that of other passerines like, the Grey Wagtail and Eurasian Blackcap.

### 13. Large axons outline the magnified foveal representation within the optic tectum

Ganglion cell axons from the temporal and central foveae project, respectively, into the MT1-MT2 and MT3 sectors of the OT (Figure 11A; Rodrigues et al., 2023). We wondered whether the large axons originating from FMGCs localized in the temporal fovea projected to the OT or to other midbrain structures. Using serial semi-thin sections, we counted axons with a diameter ≥ 3.5 µm (cross-sectional area ≥ 10 µm²) within the stratum opticum (SO) of sectors MT1, MT2, and MT3 in the Barn Swallow Hr21 (Figure 11B, C). In the sectors MT2 and MT3, sections were basically parallel to the rostro-caudal axis, whereas they were perpendicular in MT1 (Figure 11A). We have counted a total of 560 axons ≥ 3.5 µm across the entire surface of the SO. They were preferentially localized in the MT2 sector. The fact that the total count was in the range of that obtained in the ON (487 axons > 10 µm^2^; Figure 10B), makes it likely that all the large axons that originated from the FMGCs project onto the OT. The slightly higher number of axons counted in the SO compared to the ON can be explained by the fact that for some axons, the cutting plane was perpendicular or oblique to their trajectories and might have been counted twice on serial sections. Although the area covered by large axons in the OT was more difficult to measure than in the ON, we estimated that large axons were distributed over an area of approximatively 2 mm^2^ while the total surface area of the SO was estimated between 12 and 15 mm^2^ (Figure 11D). This result contrasted with the grouping of foveal GCs ≥ 200 µm^2^over an area of only 0.7 mm^2^ in retinas whose surface areas were estimated at 55-60 mm^2^. The broad distribution of large foveal axons in the SO may reflect magnification of the temporal fovea within the OT. In the House Martin, although large axons were less numerous than in the Barn Swallow (Figure 10B), they were still distributed in a broad but discrete area of the SO (Figure S10).

**FIGURE 11.**
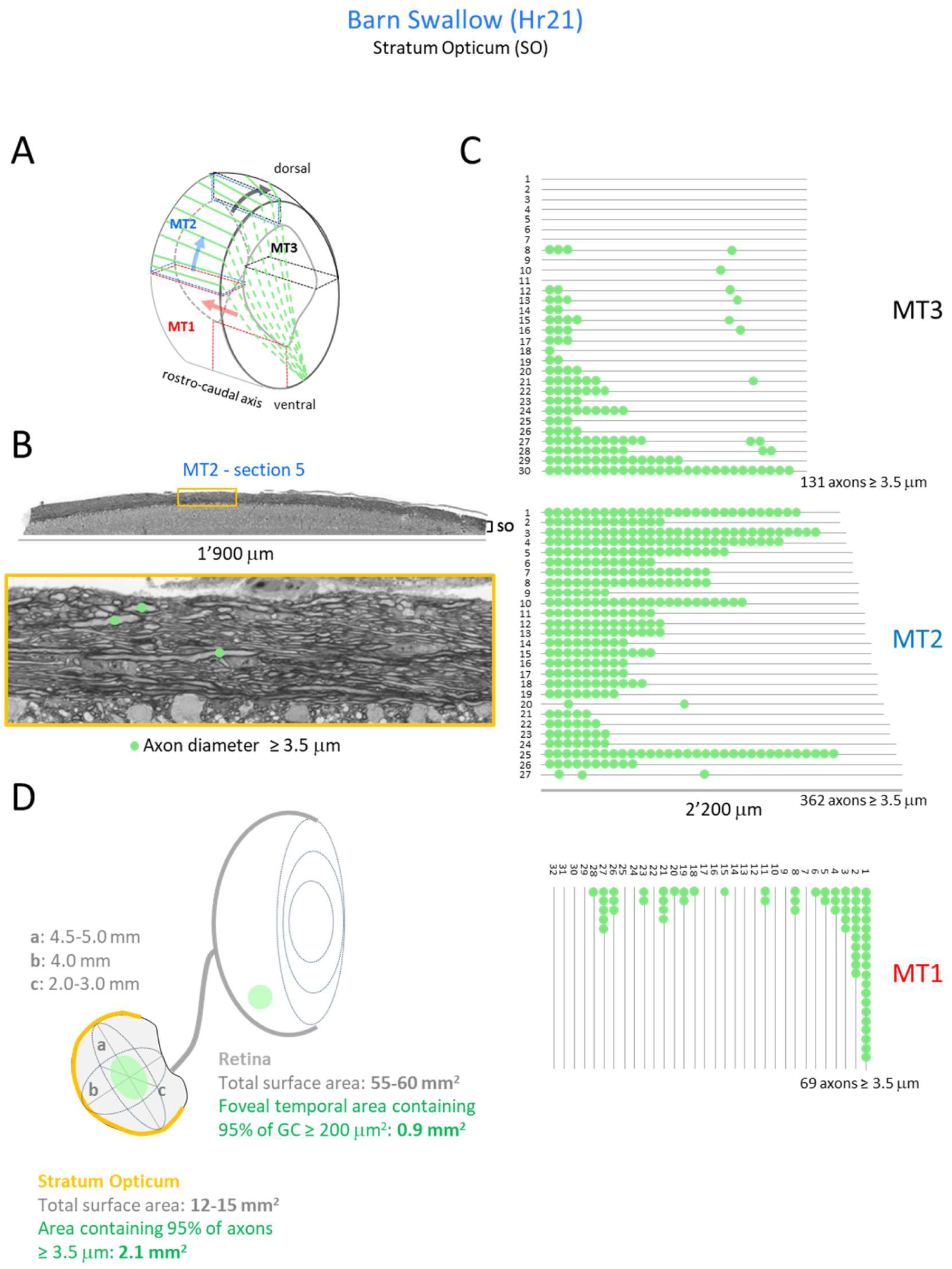
Topographic distribution of large-diameter axons (≥ 3.5 µm) in the SO of the Barn Swallow. (**A**) Selection of three medio-temporal sectors (MT1, MT2, MT3) of the optic tectum (OT) for axon counting. Rectangles and arrows indicate cutting planes and directions, respectively; green lines schematically represent axonal organization (**B**) Semi-thin section of the SO in the MT2 sector. Inset: Axon diameters are measured at the point of maximum thickness along the longitudinal plane. (**C**) Axon counts (≥ 3.5 µm) performed on serial sections at 30 µm intervals. There is a continuum between the sector MT2 and MT3 and the section 30 of MT3 is at less than 50 μm from the section 1 of the sector MT2. (**D**) Schematic representation comparing areas occupied by big GC (≥ 200 µm²) in the temporal fovea and large axons (≥ 3.5 µm) in the SO.

**FIGURE S10 (related to FIGURE 11).**
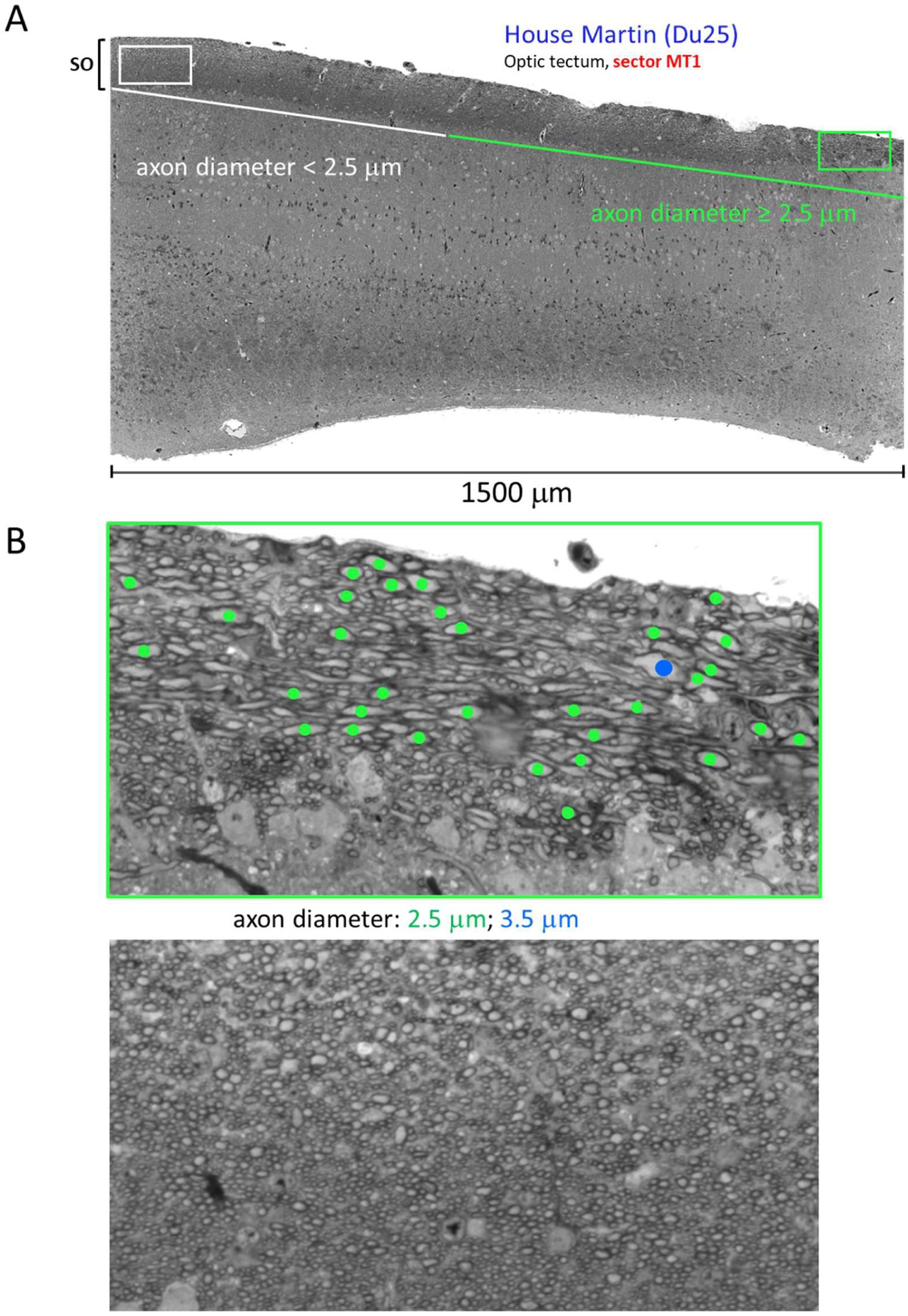
Distribution of large-diameter axons in the SO of the House Martin. (**A**) Large axons (≥ 2.5 µm in diameter) are selectively distributed within sector MT1, appearing in a broad but discrete region (delineated by the green line), while absent in sectors MT2 and MT3. (**B**) High-magnification views of the SO showing a region containing large-diameter axons (green frame) and a region devoid of such fibers (white frame). Colored dots represent axon diameters: green = 2.5 µm; blue = 3.5 µm.

## DISCUSSION

Our study reveals the primary role the fovea would play in motion detection. The large axons originating from a population of foveal GCs with somas ≥ 200 μm^2^ delineate the area occupied by foveal axons within the ON and the extent of foveal magnification within the OT. The primary difference in retinal organization between aerial insectivorous birds and other bird species lies in the organization of their temporal retinas. In the Barn Swallow, the House Martin and the Common Swift, a deep temporal foveal pit delimits two different areas. Fovea asymmetry that arises from changes in the spatial distribution of cone and GC subtypes enable swallows and swifts to perform the same task, i.e., identifying and catching one insect every 10 to 20 seconds while flying at speeds of 40 to 60 km/h. Slonaker (1897) and Polyak (1957) did propose that in swallows, the central retina is primarily responsible for the lateral and monocular vision, while the temporal retina is used for frontal and binocular vision. Martin (2022) put forward the idea that in birds, frontal vision is concerned with the detection of movement, but not with the detection of detail, and high spatial resolution is in the lateral fields and used to detect objects. Comparative analysis of bird retinas (Rodrigues et al., 2023) and data presented in this study provide us with a more nuanced view. For instance, in the Common Swift and the Barn Swallow central retinas, about 30 cones pool their signals onto a single GC, suggesting that, at least in these species, lateral vision is well adapted to detect moving objects. Furthermore, in these species, the temporal foveae are the retinal areas with the lowest convergence of cones onto GCs, suggesting that they are the most appropriate for detecting details of objects as they become visible from a distance in the frontal visual field. Polyak (1957) noted that when an object moves rapidly in the front of a Barn Swallow’s beak at a distance of 15 to 20 centimeters, both eyes converge upon it and follow it. At this short distance, according to our data, it is the temporal foveal area oriented towards the periphery that is best suited to track the object. The thick cones with short outer segments present in this area are indeed adequate for detecting nearby objects due to their potentially wider receptive fields and lower photopic light sensitivity as compared to the thin cones with long outer segments lying across the foveal pit. When an object comes into view in the frontal visual field and the signal crosses the temporal fovea, the bird might have a few milliseconds to adjust and stabilize its flight for managing the interception. Neurons responsible for detecting movement confront a phenomenon known as the *aperture problem* (Wuerger et al., 1996), especially when detecting movement of nearby objects. Indeed, a neuron cannot detect the direction of an object moving across the visual field when the object is larger than its receptive field. The big foveal GCs and their putative wide dendritic trees can contribute to overcome this problem by providing broader receptive fields. The sizes of their somas localized in the GCL place them in the category of the largest GCs identified in birds (Chen and Naito, 1999, 2009; Coimbra et al., 2014; Coimbra et al., 2009; Hayes et al., 1991). However, they distinguish themselves from other big GCs by their quasi-exclusive localization in the temporal fovea. In the absence of biochemical or electrophysiological specification, it remains unclear whether they compose a discrete cell type. They are nevertheless good candidates for motion detection and we named them *foveal magnocellular-like ganglion cells* (FMGC). FMGCs are abundant in the temporal foveae of the swifts and swallows, whereas they are rare or absent in the temporal retinas of birds - like the Common Kestrel, the Eurasian Sparrowhawk and the Grey Wagtail - that use a perch-and-sally feeding technique. Identifying, tracking and catching moving prey on the wing is a more complex task than doing so from a stationary, fixed frame of reference. FMGCs would be involved in achieving this complex task. In their pioneering study, Maturana and Frenk (1963) have shown in the pigeon retina that an individual GC identified with the help of an electrode placed in the optic nerve can respond to a bar moving vertically (upward and downward) but not horizontally across its receptive field. This and Levick (1967) study on the rabbit mark the discovery of orientation-selective ganglion cells (OSGC). Since then, studies on retinal orientation selectivity have focused almost exclusively on rabbits and cats and more recently on mice (reviewed in Antinucci and Hindges, 2018). Direction-selective retinal ganglion cells and circuits for direction selectivity have now been identified in primates (Kim et al., 2022; Patterson et al., 2022; Wang et al., 2023). On the other hand, no GC type exclusive to the primate fovea has yet been found. In House Martin, FMGCs are smaller and less numerous than in the Barn Swallow and the Common Swift and would not have been detected without analyzing the ON. In the Barn Swallow, we propose a model where the about 400 FMGCs are distributed in 80 clusters over an area of 0.7 mm^2^, each made up of 5 FMGCs, (Figure 12). We postulate that the FMGCs of one cluster have overlapping dendrites and receive their inputs from approximatively 100 cones. Each FMGC may have an excitatory region parallel to an inhibitory region, much like a OSGC or a cortical simple cell of the cat or monkey (Hubel and Wiesel, 1962). This situation is plausible in view of a study in mice (Nath and Schwartz, 2017) showing that two OSGC with neighboring somata and overlapping dendrites exhibit orientation-selective responses and that their preferred orientations are different. In our model, we suggest that depending on its orientation and position, the moving object may preferentially activate one of the 5 FMGC of clusters overlaying the foveal area. These FMGCs should enable the bird to adjust and stabilize its flight in order to keep the image of a selected moving object in a functional focus area. The bird has a few milliseconds to identify the object and decide whether to catch it (e.g. a harmless insect) or avoid it (e.g., a harmful obstacle). In this context, we can ask how the fast-flying bird distinguishes between a moving object and a self-induced moving background, i.e., the optic flow field. Experiments with pigeons and hummingbirds have revealed that the tectofugal system is well-adapted to respond to various aspects of object motion while ignoring self-induced visual motion. Conversely, the accessory optic system is well suited to extract important features of self-induced visual motion (Frost et al., 1990; Gutierrez-Ibanez et al., 2025). The fact that the large axons originating from the FMGCs innervate the OT rather than elements of the thalamic or accessory pathways suggests that the swifts and swallows might be better at distinguishing a rapidly moving object from its context than pigeons. Indeed, pigeons utilize their non-foveate temporal retinas, i.e., the red fields, for tasks requiring fine detail perception and binocular vision, such as grasping seeds, rather than for detecting motion in their frontal field (Querubin et al., 2009). In birds that feed on the ground, a lesion of the isthmo-optic nucleus (ION) suppresses centrifugal axons that innervate the retina and impair the bird’s capacity at distinguishing a specific object, e.g., a seed, from a distracting background (Uchiyama et al., 2012). In the Common Swift and Barn Swallow, there are fewer neurons in the ION than in ground feeding birds, suggesting that aerial birds are differently adapted to focus on a specific object (Feyerabend et al., 1994; Wilson and Lindstrom, 2011).

**FIGURE 12.**
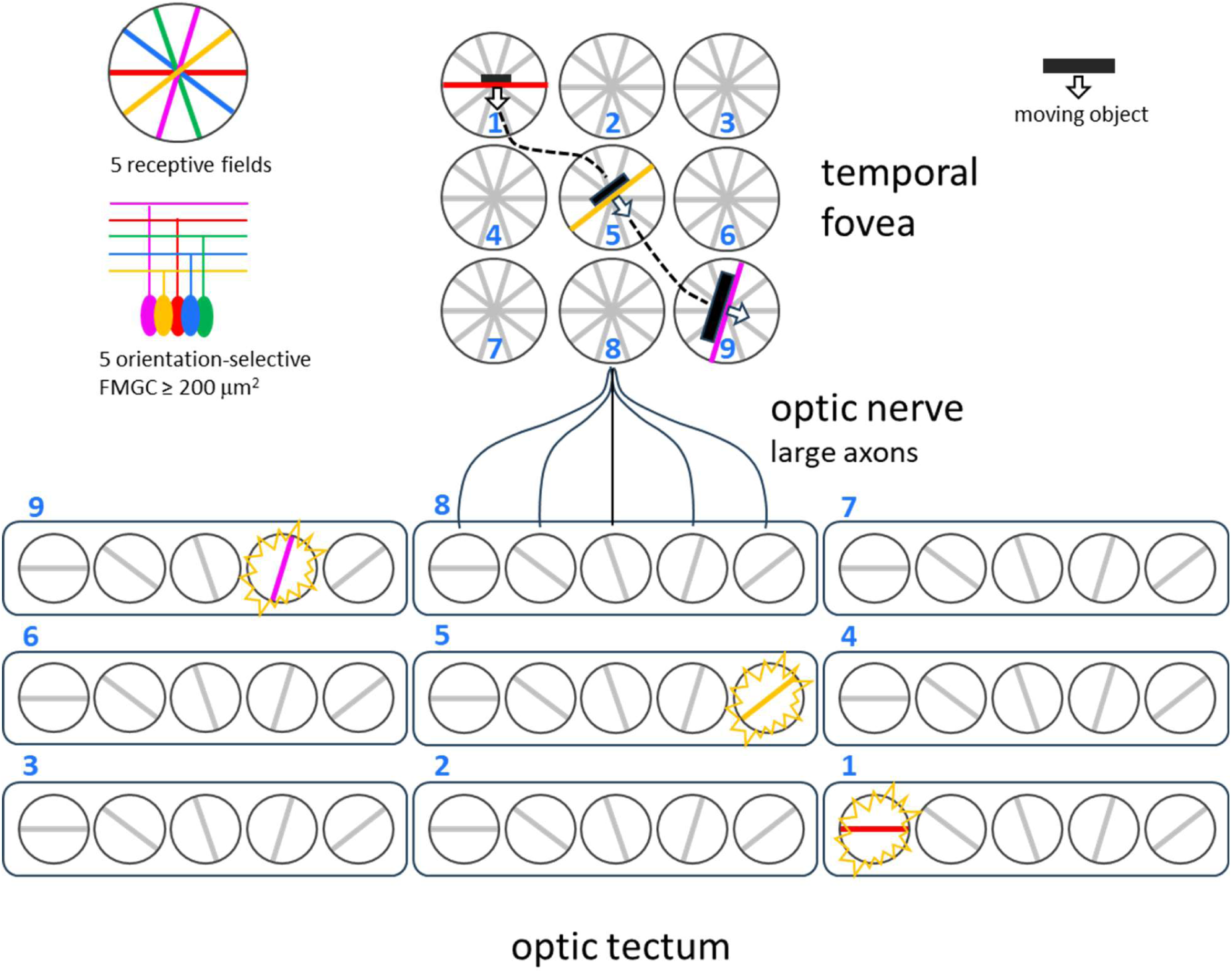
Proposed model of adversarial dialogue between FMGCs and midget-like GCs in the avian fovea. Tiling of the foveal area by motion-detecting ganglion cells (FMGCs) and midget-like GCs, providing the neural substrate for integrated spatial and temporal acuity. In the Barn Swallow and Common Swift, ∼400–600 FMGCs (∼4% of total foveal GCs) are organized into 80 clusters surrounding the foveal pit over an area of 0.7–1.0 mm². Each cluster is modeled with a ratio of 5 FMGCs, 100 midget-like GCs, and 200 cones. Functional roles: Midget-like GCs mediate object identification in the frontal visual field, while FMGCs modulate flight trajectory via feedback loops with the optic tectum to maintain the target within their receptive fields. Overlapping receptive fields within each cluster ensure orientation-independent detection. Adversarial interaction: FMGC-mediated suppression of midget-like GC responses enhances selective attention through target/background progressive discrimination during pursuit, preventing target switching and maximizing capture efficiency. The dispersed connectivity of FMGCs in the optic tectum reflects foveal magnification.

Large axons originating from FMGCs localized in the temporal foveae of the Barn Swallow, the House Martin and the Common Swift occupy well delimited areas in the ON that are proportionally larger than the surface areas occupied by the temporal foveae in the retinas. This pattern may prefigure the dispersion of large axons in the stratum opticum of the OT. The broad representation of the temporal fovea in the ON and in the OT occurred despite the fact that GCs localized in the temporal fovea represent only 4% of all GCs in the Barn Swallow and 3% in the Common Swift (estimated from the total number of axons in the ON). Clarke and Whitteridge (1976) have used electrophysiological techniques to identify a magnified representation of the central fovea and of the red field within the pigeon OT. They concluded that the OT allocates a larger area to the fovea and red field compared to other regions of the retina because of their higher densities of GCs. However, at that time, the authors were not aware that the pigeon adapts to the increase of retinal inputs by increasing the density of tectal neurons (Rodrigues et al., 2023). In the Barn Swallow, we suppose that the distribution of 560 axons ≥ 3.5 μm in diameter over a well-delimited area covering ∼16% of the total surface area of the stratum opticum, outlines the foveal representation within the OT. The question remains as to why a few hundred large axons are distributed over such a wide tectal area. One reason could be that their broad axon terminals synapse with a greater number of tectal neurons. The number and density of tectal neurons in the MT2 sector are indeed higher in the Common Swift the Barn Swallow and the House Martin than in other bird species, despite lower densities of GC in their temporal retinas (Rodrigues et al., 2023). Congruent with this idea, Azzopardi and Cowey (1993) have shown in the rhesus monkey (*Macaca mulatta*) that the cortical representation of the fovea is greater than expected from the density of foveal GCs, allowing them to conclude that there (…) *is more cortex per GC in the fovea than there is on average in the entire projection* (Azzopardi and Cowey, 1996).

In summary, the identification of putative, motion-sensitive foveal ganglion cells (FMGCs) interspersed among midget-like circuits, and their involvement in foveal magnification within the brain, challenges established models of visual processing by suggesting the integration of motion sensitivity at the foveal level. Elucidating the exact structural morphology of these FMGCs and how they interact with ACs and midget-like pathways will be essential for understanding how the visual system seamlessly integrates spatial and temporal acuity.

## METHODS

Our core strategy involved cross-species comparison of the visual systems of 10 bird species that belong to 7 families. In each of the 19 specimens analyzed (Table S1), 12 tissue samples were collected from the retinas, the optic nerves and the midbrain. The 228 tissue samples were processed according to a standardized protocol ensuring consistent and high-quality histology on semi-thin plastic sections. We have developed custom tools for analyzing microscopy images.

### Animals

The specimens used in this study are wild birds that suffered from unrecoverable injuries (Table S1) and had to be euthanized at the rehabilitation centre La Vaux-Lierre (http://www.vaux-lierre.ch) in accordance with Federal Swiss veterinary regulations. The decision to euthanize made by one of us (EB) was independent of the project. Adult Japanese Quails were provided by Marc Plancherel (https://fribourg.ch/fr/terroir-fribourg/producteurs/oeufs-de-cailles-marc-plancherel).

### Tissue isolation and processing

The retinas, the ON and the OT were dissected in DPBS (ThermoFisher). Tissue were prefixed in 3.7% formaldehyde, 1% glutaraldehyde, 0.1 M Sodium phosphate monobasic, 0.07 M NaOH in distilled water at 4°C for 30 min. To prevent tissue degradation, the initial fixation must occur within 20 min of confirmed cardiac death. Beyond this window, cellular decay becomes evident during histological analysis of semi-thin sections, with FMGCs being particularly susceptible to these post-mortem changes. Initial fixation stiffened the tissue for precise cutting. We then incubated 2–4 mm cubes of retina and OT, plus 3–4 mm segments of the proximal ON, at 4°C until embedding. The tissue samples were washed with 0.1 M ice cold sodium cacodylate before incubation 10 min in 0.8% K_3_Fe(CN)_6_ in 0.1 M sodium cacodylate and fixation in 1% osmiumtetroxide, 0.8% K_3_Fe(CN)_6_ in 0.1 M sodium cacodylate pH 7.4 for 90 min. They were washed with sodium cacodylate and water, stained 2 hours with 1% uranyl acetate, dehydrated in ethanol baths and incubated in propylene oxide 100% for 45 min, followed by propylene oxide/epon 1:1 incubation overnight. Tissue cubes were embedded in the proper orientation in epoxy resin consisting of 48% Agar 100, 18% DDSA, 31% MNA and 3% BDMA (agar scientific). Semi-thin (1 µm thick) sections were produced using a Leica UCT ultra-microtome and were mounted on glass slides (SuperFrost/plus, Assistant).

### Quantifying cell density and cellular area within retinal and tectal cross-sections

Plastic sections 1 µm thick from retina and OT mounted on glass slide were stained by incubation 1-2 minutes in 1% toluidine blue, 1% sodium borate in distilled water. Coverslips were sealed with xylene-free EUKITT® neo (O. Kindler ORSAtec). Images were acquired with a Zeiss LSM 800 confocal microscope. Using tile region acquisition, we imaged areas with dimensions that exceeded the size of an individual image field. Tiles were stitched together using Zen, Zeiss image-processing software. To count the number of cells in retina and axons in the Stratum Opticum of the OT, each cell body and axon was manually annotated by surrounding it with a closed line using an image analysis software (ImageJ/Fiji).

### Computing axon number and axon area on ON sections

#### Imaging ON axons

The best-preserved ON tissues were identified less than 20 µm from the block surface. Analysis was restricted to 1-µm-thick cross-sections that exhibited homogeneous 1% toluidine blue staining and were free of vacuolated areas. Using a Zeiss LSM 800 confocal system, we acquired and stitched image tiles with ZEN software. This process ensured that even the smallest axons remained sharp across the full surface of the optic nerve section.

#### Morphometric analysis of the ON

##### 1. Automated detection of axons

We trained and used a deep learning algorithm for image segmentation to automatically detect and mark axons in all images of ON sections that were used in this study.

##### 2. Algorithm

We used an existing Python implementation based on the Keras framework of a convolutional neural network (CNN) called U-Net (Ronneberger et al., 2015). As argued by the authors, this neural network architecture combined with data augmentation techniques, has proven to be efficient for cell detection in biomedical image data, especially when few labeled data is available.

We started from an existing implementation based on Python’s Tensorflow and Keras frameworks. The original implementation is available online (https://github.com/zhixuhao/unet) under a MIT License. The package versions we used are: Tensorflow (v2.13.0) and Keras (v2.13.1). The exact implementation of the CNN algorithm that we used in this study is available at https://github.com/bird-visual-system/onunet.

##### 3. Training data

We trained the network from scratch with manually labeled data subsampled from ON images of three species: Barn Swallow, Common Swift and Domestic Rock Pigeon. Each image was subdivided into 512 x 512 pixels tiles. A typical 512 x 512 tile contains around 1000 axons. Then, from a set of tiles which were randomly selected among the three samples, 34 tiles were selected and manually annotated using an image analysis software (ImageJ/Fiji) by precisely surrounding each axon. From that, binary images were generated, with white pixels corresponding to axons and black to anything else (background). The set of labeled tiles was then randomly splitted into two subsets: one for training (80%) and the remaining for validation (20%).

In order to increase the size of the training set, we used standard data augmentation techniques, by performing elastic deformations, rescalings, reflections and rotations of the training data.

##### 4. Training setup

As suggested in Ronneberger et al. (2015) in order to improve boundary detection, we used a weighted loss where pixels separating neighboring axons obtain larger weights than others. We then used these weights in a weighted binary cross-entropy loss function as explained below.

Let us denote by *X* a tile from the original image, let *Y* denote the binary (black and white) image obtained from the manual labeling and let *Ŷ* denote a prediction issued by the algorithm. In our setting, *X*, *Y*, and *Ŷ* are 2-dimensional tensors (matrices) of size 512 x 512. An element *x_ij_* of *X*, 0 ≤ *x_ij_* ≤ 1, represents the value of a pixel of the original grayscale image. An element *y_ij_* ∈ {0,1} of *Y* represents the value of a pixel of the binary label: *y_ij_* = 1 (white) means that pixel (*i*, *j*) belongs to an axon, whereas *y_ij_* = 0 (black) means that pixel (*i*, *j*) belongs to anything else (background). An element *p_ij_*, 0 ≤ *p_ij_* ≤ 1, of the prediction *Ŷ* issued by the algorithm is the probability that pixel (*i*, *j*) belongs actually to an axon.

During the training phase, the algorithm learns to issue predictions *Ŷ* in order to minimize a loss function. In our case, the loss function is defined as

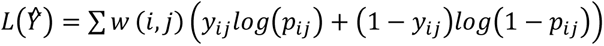

where the weight function *w*(*i*, *j*) is given by

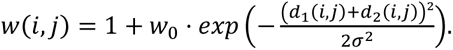

The purpose of the weight function resides in giving more weight in the overall loss *L*(*Ŷ*) to pixels separating two axons so that the algorithm pay more attention during training to correctly predicting them. In the latter expression, *d*_1_(*i*, *j*) denotes the distance of pixel (*i*, *j*) to the closest axon, *d*_2_(*i*, *j*) denotes its distance to the second closest axon, whereas *w*_o_ and *σ* are fine-tuning parameters. After experimenting with different values, we set *w*_o_ = 3 and *σ* = 4. The weight function *w*(*i*, *j*) was used to precompute the weights of the pixels in each of the manually labeled images used for training, providing thus for each training/validation image a weight matrix *W* of the same dimensions than *X* and *Y*.

The neural network was then trained using “Adam stochastic gradient descent” algorithm implemented in Keras, during 100 epochs and 200 steps per epoch.

##### 5. Making predictions

Once the training completed, the neural network was used to make predictions on each of the 512 x 512 tiles forming the partition of a whole ON image. The predicted tiles were then stitched together to form an almost segmented image of the whole ON in which pixels that are likely to belong to axons are (close to) white, whereas pixels that are likely to not belong to axons are (close to) black.

##### 6. Further analysis

The grayscale image of the whole optic nerve was then converted to a binary image using a experimentally set threshold of 0.7 to achieve a good compromise between border detection and axon area underestimation. Then, a standard contour detection algorithm (Python’s *skimage.measure.find_contours* implementation of the “Marching squares” algorithm) was used to extract axon borders from the predicted binary mask of the whole optic nerve.

Morphometric analysis was performed in Python with the help of the following packages: scikit-image (v0.21.0) and shapely (v2.0.1) for contours extraction and geometrical analysis of the axons, seaborn (v0.12.2) for the spatial statistical visualization (heatmaps), matplotlib (v3.7.2) for graphs, numpy (v1.24.3) for scientific computing, pandas (v2.1.0) for data processing. All scripts used for the analysis are available at https://github.com/bird-visual-system/cellcount.

### Analysis of the ON by transmission electron microscopy (TEM)

The ON were embedded in epoxy resin and 80–90 nm sections were produced using a Leica UCT ultra-microtome, and were recovered on TEM grids. Staining was done by incubation 5 min in Acetone/Uranyl Acetate 1:1 solution followed by 10 min in lead citrate solution. TEM images were acquired using a Tecnai G2 Transmission Electron Microscope. To compute the number and area of optic nerve axons, each myelinated axon was manually annotated by surrounding it with a closed line using an image analysis software (ImageJ/Fiji).

## ACKNOWLEDGMENTS

We thank Laurent Brodier for his expertise in preparing optic nerve samples for Transmission Electron Microscopy. Tissue processing and confocal microscopy were performed at the Bioimaging Platform of the Faculty of Sciences (http://bioimaging.unige.ch) and we wish to thank Christof Bauer and Jérôme Bosset for their advice. We are grateful to Damien Juat for having organized a fruitful meeting with BG who then joined the project. We are grateful to Marc Ballivet (University of Geneva) for the critical reading of the manuscript. Special thanks to Laurent Vallotton (Natural History Museum of Geneva) for his ornithological advice.

